# On biological networks capable of robust adaptation in the presence of uncertainties: A systems-theoretic approach

**DOI:** 10.1101/2022.09.23.509157

**Authors:** Priyan Bhattacharya, Karthik Raman, Arun K. Tangirala

## Abstract

Biological adaptation, the tendency of every living organism to regulate its essential activities in environmental fluctuations, is a well-studied functionality in systems and synthetic biology. In this work, we present a generic methodology inspired by systems theory to discover the design principles for robust adaptation, perfect and imperfect, in two different contexts: (1) in the presence of deterministic external disturbance and (2) in a stochastic setting. In all the cases, firstly, we translate the necessary qualitative conditions for adaptation to mathematical constraints using the language of systems theory, which we then map back as design requirements for the underlying networks. Thus, contrary to the existing approaches, the proposed methodologies provide an exhaustive set of admissible network structures without resorting to computationally burdensome brute-force techniques. Further, the proposed frameworks do not assume prior knowledge about the particular rate kinetics, thereby validating the conclusions for a large class of biological networks. In the deterministic setting, we show that unlike the incoherent feed-forward network structures (IFFLP), the modules containing negative feedback with buffer action (NFBLB) are robust to parametric fluctuations when a specific part of the network is assumed to remain unaffected. To this end, we propose a sufficient condition for imperfect adaptation and show that adding negative feedback in an IFFLP topology improves the robustness concerning parametric fluctuations. Further, we propose a stricter set of necessary conditions for imperfect adaptation. Turning to the stochastic scenario, we adopt a Wiener-Kolmogorov filter strategy to tune the parameters of a given network structure towards minimum output variance. We show that both NFBLB and IFFLP can be used as a reduced order W-K filter. Further, we define the notion of nearest neighboring motifs to compare the output variances across different network structures. We argue that the NFBLB achieves adaptation at the cost of a variance higher than its nearest neighboring motifs whereas the IFFLP topology produces locally minimum variance while compared with its nearest neighboring motifs. We present numerical simulations to support the theoretical results. Overall, our results present a generic, systematic, and robust framework for advancing the understanding of complex biological networks.

## 1 Introduction

The evolutionary drive for survival requires every living organism to perform several essential functions that emerge from the underlying complex biological systems [1]. Adaptation, a ubiquitous property in survival mechanisms, refers to the ability to sense the change in the external disturbance and revert to it’s pre-disturbed operating conditions [2–4]. Due to the regulation aspect, adaptation plays a crucial role in a variety of phenomena ranging from bacterial chemotaxis, mammalian Ca^2+^ homeostasis to temperature regulation in the human body [5–10]. Therefore, it is customary to study the emergence of perfect adaptation in a way that not only aids in the understanding of the constituents of the biological process underneath adaptation but also is convenient for synthetic design and therapeutics— this necessitates an intervention inspired by systems biology, for it offers equal importance to both the constituents of a biochemical process and their interconnections.

According to the formalism of systems biology, a biological functionality can be thought of as a response emanating from a complex network [11]. Given the knowledge about the constituents of networks from classical disciplines of biology, it is an important task to establish a mapping between the output response and the architecture of the network (*design principles*) [2]. Further, it is widely believed that the design principles for a given functionality are conserved across the organism space [3, 4]. For instance, adaptation required bacterial chemotaxis of *E. coli* is obtained by negative feedback between two protein complexes, whereas adaptation involved in regulating Ca^2+^ level in mammalian cells also necessitates negative feedback between the parathyroid hormone and vitamin D [9]. Despite the structural similarity, it is not necessary for the underlying rate dynamics to be identical in both the cases [12]. Therefore from the perspective of theoretical investigation, it is expected that any methodology designed to identify the network structures for any given functionality should not be confined to a specific rate dynamics [13].

Previous approaches to deducing the design principles for different adaptation variants can be broadly classified into three categories: (i) computational, (ii) rule-based, and (iii) systems-theoretic [14]. Typically, a biochemical network is represented by nodes being the biochemical species and the edges symbolizing the chemical reactions. Each edge can be of two types, activation and repression, depending on the chemical reaction [4]. Therefore, considering the directionality and type of each edge, an N–node network can have 3^N^2^^ possible network structures [4]. Further, each network structure entails a system of differential equations constituted by the chemical rate equations for each reaction. The computational approach involves examining all possible network structures via simulating the underlying dynamical system of each structure. Furthermore, each network structure is simulated by multiple sets of rate constants *i. e.* parameters. Ma *et al* (2009) implemented a computational approach to identify 3–node network structures that can provide perfect adaptation in the presence of step-type disturbance [4]. The performance of each network, in the context of adaptation, was measured by two parameters, namely, *sensitivity* (S) and *precision* (P). As it can be observed from Fig. 2, sensitivity and precision should be non-zero finite and infinity respectively in the case of perfect adaptation. With an assumption of Michaelis–Menten rate kinetics, it was observed that each network structure capable of providing adaptation (S ⩾ 1, P ⩾ 10) contains either negative feedback with buffer action (*NFBLB*) or incoherency between the forward paths from the input-receiving node to the output node (*IFFLP*) [4].

**Fig 1.**
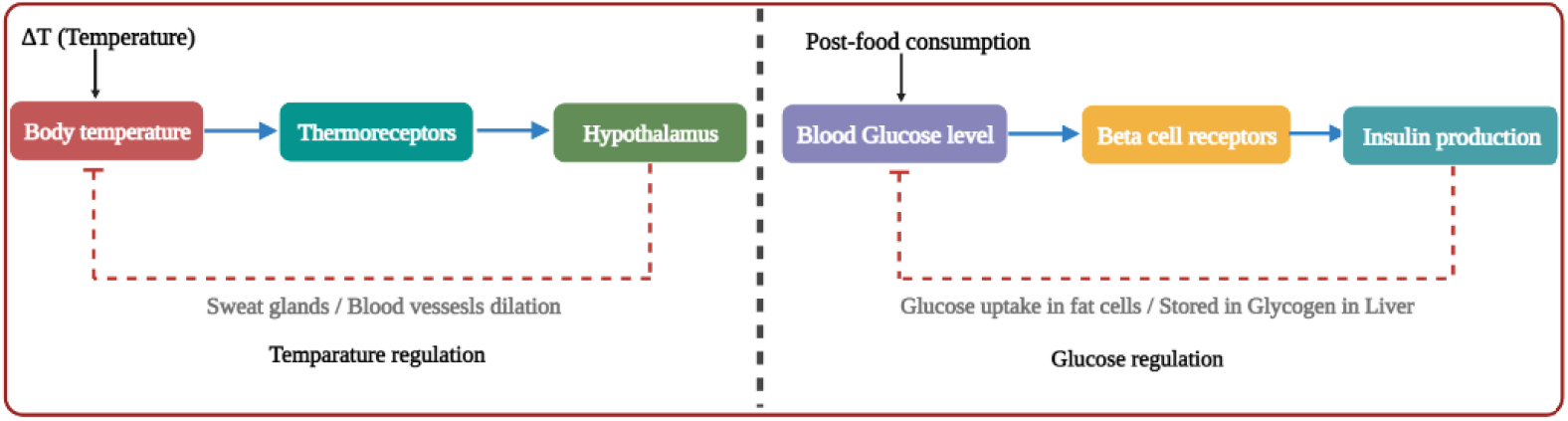
Conservation of design principles. The figure on the left represents an abstraction of the temperature regulation process whereas the figure on the right depicts the glucose regulation process of mammals. Despite the difference in the network constituents, both these processes involve a negative feedback loop to achieve perfect adaptation. The *hypothalamus* and the Insulin production markers serve as the controller species for temperature and glucose regulation processes respectively.

**Fig 2.**
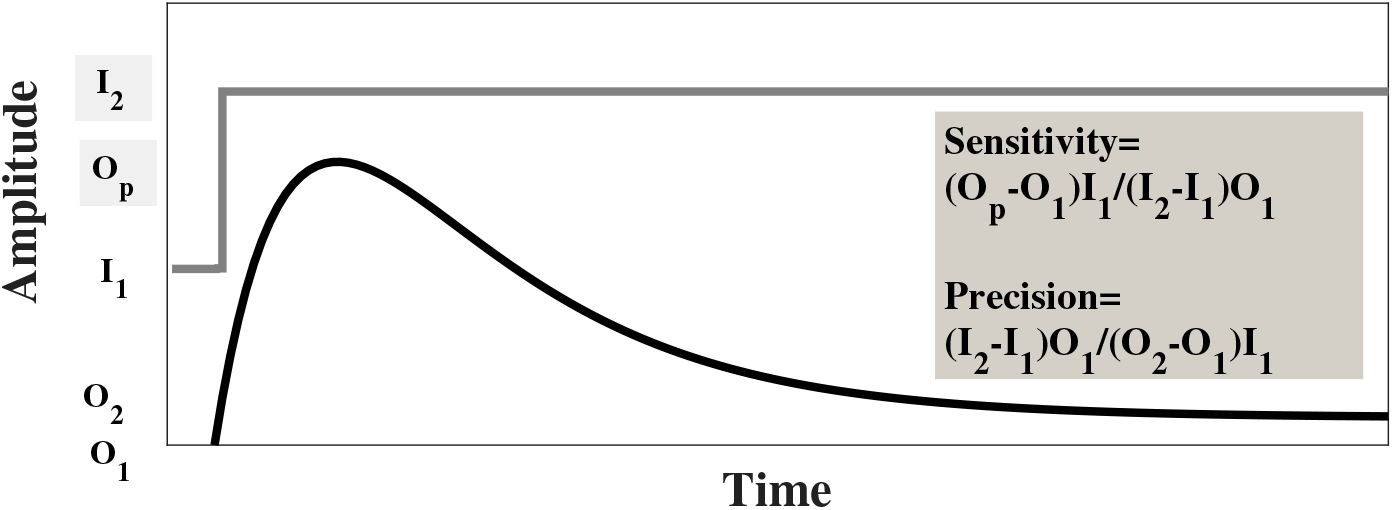
A typical response for adaptation

Further, the ratio between the number of parameter sets for which a given network structure exhibited adaptation to the total number of simulations performed for that structure is considered an empirical measure (Q) of robustness. It was found that an NFBLB yields better performance than IFFLP in terms of robustness, and the combination of IFFLP and negative feedback can improve the robustness of IFFLP for a 3-node network. Later, Qiao *et al* (2019) extended this approach to non-deterministic disturbances [15]. The input disturbance, in this case, was considered to be synthesized from a chemical Langevin’s equation. The ratio between steady-state output and input variance was also measured with perfect adaptation as regulation of the expected value of the output at steady-state. Interestingly, it was found that there exists a trade-off between sensitivity and the noise ratio for a class of adaptive networks. In contrast, simultaneous achievement of infinite precision and noise attenuation was considered viable. Further, Otero-Muras and Banga (2019) implemented a mixed-integer nonlinear optimization program to deduce the network structures for adaptation [16].

The rule-based methodology relies on the existing engineering strategies to design biological networks for the desired requirements. Briat *et al* (2016) provided an antithetic feedback control strategy to design a biological network capable of adaptation. It was shown that the antithetic integral control provides perfect adaptation even in parametric uncertainties [17]. It was assumed that the sensor reactions for the proposed controller module are not disturbed by parametric fluctuations. Later, it was observed that the antithetic integral controller performs perfect adaptation at the cost of increased variance—a phenomenon to be regarded as the stochastic counterpart of the well-known destabilizing tendency of a pure integral controller in the deterministic domain [18]. Therefore, contrary to the findings by Qiao *et al* (2019), Khammash *et al* (2018) suggested a trade-off between *precision* and the *output variance* for the antithetic feedback controllers. As a remedy, additional negative feedback has been proposed for some range of parameters that render the output variance lower than that in the open-loop scenario.

Unlike the previous two approaches, the systems-theoretic approaches begins with characterizing the functionality through several performance indices in such a way that there exists a well-defined mapping between these indices and a few of the standard parameters relevant to the systems-theoretic analysis. Therefore, the performance indices evaluated for the ideal scenario (*i.e.* in the case of best possible performance) translates to several *precise mathematical equations* involving the systems-theoretic parameters. Further, the admissible topologies are obtained by exploiting those mathematical equations with specific assumptions about the network structure and the corresponding dynamical system. Systems-theoretic contributions to understanding adaptive biological networks can be divided into two categories—the first set of approaches primarily aimed at identifying the structure for **perfect adaptation**. In contrast, the second set focused on the robustness of the system and imperfect adaptation. Sontag (2003) first argued that perfect adaptation is a well-known regulation problem in control theory [19]. Therefore, adaptation in the presence of a step-type disturbance requires an integrator within the system. Using this result, Allowgner *et al* (2012), Drenstig *et al* (2008), Bhattacharya *et al* (2018) obtained the condition for perfect adaptation in terms of the systems theoretic parameters [20–23]. Further, Bhattacharya *et al* (2018) obtained the necessary structural requirements for perfect adaptation in a network of three nodes [23]. Additionally, a motif-specific optimization problem was also suggested to obtain the desired set of parameters (rate constants) for the structurally admissible (for adaptation) topologies. Later Araujo *et al* (2018) and Wang *et al* (2020,2021) adopted a graph-theoretic method to unravel the network structures for perfect adaptation in a network of any size [24–26]. It was argued that a network of any size must contain either a feedback loop facilitating the buffer action (Balancer module) or multiple forward paths with mutually opposite effects (Opposer module) to exhibit perfect adaptation. Further, Araujo *et al* suggested that the balancer module must contain negative feedback for stability [24]. Recently, we (2022) proved that a balancer module of any size, without a single negative feedback loop, renders the system unstable, thereby proving Araujo’s conjecture [27]. Further, we (2022) argued that adaptation-capable network structures are modular *i.e.* the adaptive property is not altered when the output node is connected to a downstream network [27].

The second aspect of the contribution by systems-theoretic approaches primarily focuses on understanding the emergence of *imperfect adaptation.* In this sense, analyzing imperfect adaptation brings the theory closer to the practice for almost every adaptive response, in reality, consists of finite precision. Further, establishing theoretical bounds on parameters for imperfect adaptation can also provide a compact understanding of the robustness of a given network structure in the presence of parametric fluctuations. Systems-theoretic interventions on imperfect adaptation started with Drenstig *et al* (2008), wherein different types of adaptation were classified and defined systematically [20]. Further, Drenstig *et al* (2008) translated the condition for imperfect adaptation as a requirement of at least one zero in the underlying dynamical system of the network [20]. Later, Bhattacharya *et al* (2021) proved that a three-node network performs imperfect adaptation as long as there exists one zero (with negative real part) that is placed before all three stable poles in the left half of the s—plane [28].

The computational approaches, as described previously, albeit producing reliable results, can not be scaled up for networks of arbitrary size. A computational approach also relies on the exact knowledge about the rate equations of the given network, rendering the associated conclusions valid for only that particular kinetics. Further, the rule-based methodologies, despite ensuring scalability, do not bring out the *all possible* network structures capable of adaptation. Therefore, it can not be utilized as a substitute for computational screen approaches, for it does not perform an exhaustive search of the entire set of structural possibilities. Although the systems-theoretic approaches have impressive performance compared to the other two approaches, interventions inspired by systems theory on imperfect adaptation and robustness are still an open problem. The necessary conditions provided by Drenstig *et al* (2008) for imperfect adaptation do not necessarily restrict the search space of network structures, for it can be shown that any biochemical network structure with N–nodes contains at least N – 2 zero. Further, it remains a task for systems theoretic interventions to evaluate the two acceptable network structures, the balancer, and the opposer modules, vis-a-vis robustness to parametric fluctuations. Additionally, understanding the behaviour of each network structure in the presence of a stochastic environment aids in a holistic understanding of robustness to both parametric fluctuations and exogenous random disturbances.

Motivated by the scopes as presented above, the paper presents a systems-theoretic approach to first analyze the network structures required for robust, perfect adaptation. Thus, a systematic evaluation of both the balancer and opposer modules in terms of robustness to parameter variations becomes possible. At the same time, the proposed control strategy also reflects an advancement in the existing literature of feedback control systems by extending the scope of internal model principle based design from dynamic output feedback to dynamic state feedback framework. Further, we show that it is *almost* impossible in practice for any network structure to provide perfect adaptation in the presence of disturbances running through the entire parameter space— this leads us to explore the prospects of imperfect adaptation. We propose a stricter (than Drenstig *et al* (2008)) set of necessary conditions for imperfect adaptation thereby resulting in a significant reduction in the search space of the feasible network structures. Additionally, we propose a sufficient condition for imperfect adaptation, which can aid in the synthetic design of biological networks capable of robust adaptation.

Further, we extend the concept of robustness to the exciting scenario of networks in the presence of noise due to several reasons ranging from stochastic external disturbances to unmodelled connections. The question of finding adaptive networks with minimum variance is addressed in two parts. Firstly, we compare the variances produced by both these adaptation-capable structures with its closest network structures thereby addressing all the dimensions of the variance minimization problem. Interestingly, we show that an opposer node produces minimum variance across it ‘closest’ (to be defined in the course of the manuscript) network structures whereas the balancer modules achieve perfect regulation while compromising on steady state output variance– these results clarify the apparent confusion in the existing literature and establishes the fact that *the relation between precision and SNR is motif specific.* In the second step, we address the problem of determining optimal parameters for minimizing variance of a given network structure. For this purpose, we adopt a Wiener–Kolmogorov filtering strategy to fine tune the parameters of a given network structure. We show that it is feasible for both opposer and balancer modules containing N–nodes to serve as a W-K filter of order less than N. Moreover, the assumptions through out the manuscript are satisfied by the the widely used reaction kinetics from mass-action, Michaelis–Menten to Hill kinetics along with the results being valid for networks of any size. Table 1 provides a consolidated representation for the contributions of the work in the light of the existing literature. The manuscript is arranged in the following order— Section 2 outlines the necessary frameworks for analyzing different variants of adaptation (robust, imperfect, and in presence of noise) in the lens of systems theory. Section 3 presents the application of the proposed frameworks to deduce novel insights about different properties of adaptive biological networks. The Discussion section consolidates these findings and puts contextualizes the results to the existing horizon of knowledge.

**Table 1.**
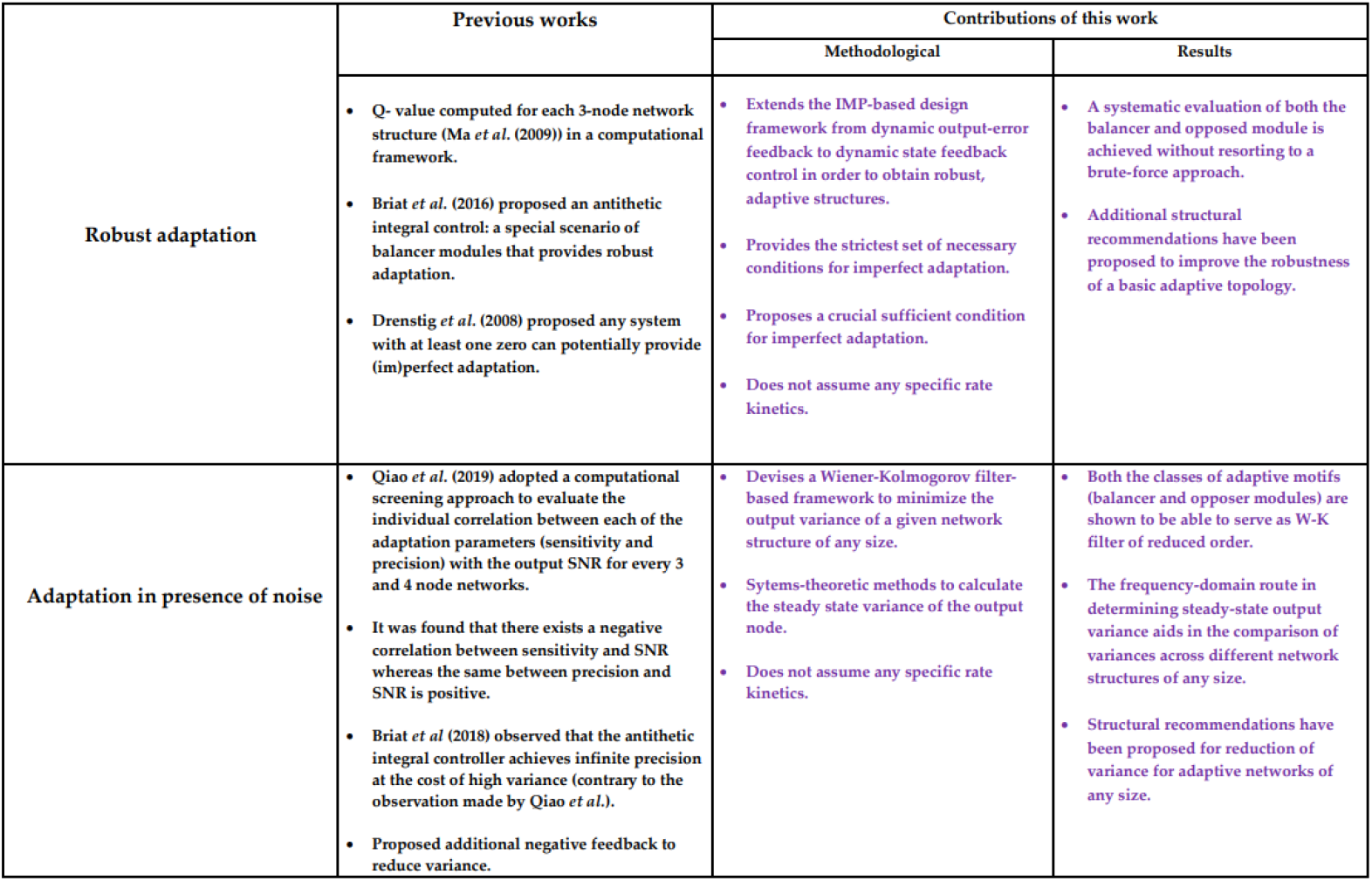
Proposed contribution of the work vis-a-vis the existing literature

## 2 Methodology

This section presents the necessary formalisms that are crucial for obtaining the main results of this work. As a first step, we propose a framework based on linear regulator theory to discover the design principles for robust, perfect adaptation. Subsequently, we propose a set of conditions required for identifying network structures capable of imperfect adaptation—this also provides a maximum bound on the parametric disturbance that an adaptive network structure can withstand. Further, we propose a generic framework for evaluating the performance of adaptation-capable topologies in the presence of various uncertainties.

### 2.1 Linearization

Since the primary aim of this work is to gain specific insights into the structure of the networks from the analysis in the realm of dynamical systems theory, it is beneficial to use linear systems theory for its well-established connection with graph networks. Hence, we start with a linearized representation of the underlying nonlinear dynamics.

For an N–node biochemical network, considering the node concentrations as the states 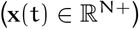, the corresponding dynamical systems can be written as

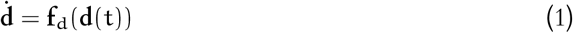

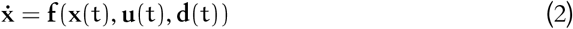

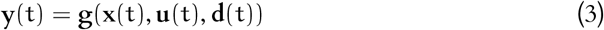

where, 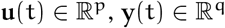, and 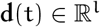 are the p inputs, q outputs and l disturbances of the system respectively. The state-space representation in the linearised domain can be written as

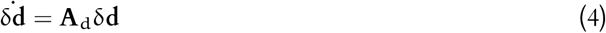

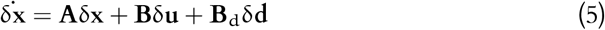

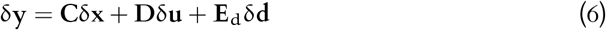

where, the matrices **A**, **B**, and **B**_d_ are obtained by evaluating Jacobian of **f** with respect to the states, inputs, and the disturbances respectively at the steady-state point. Similarly, the triplet (**C**, **D**, **E**_d_) can also be obtained by performing the same exercise **g** at steady state. Lastly, the subscript δ denotes the deviation from the point of linearization [29].

For the sake of convenience, we shall henceforth drop the δ notation from equations (5) and (6).

Evidently, the nonlinear rate functions **f** contains information about the stoichiometry and the fluxes of the system. Therefore, the Jacobian of **f** with respect to **x** serves as the *digraph* matrix for the underlying network given the following assumptions [30, 31]

1. The equations in (2) constitute a well-posed dynamical system *i. e*. **f** is locally Lipschitz with respect to **x**, **u** and **d**.
2. For a given state *x_i_* the corresponding dynamics can be written as

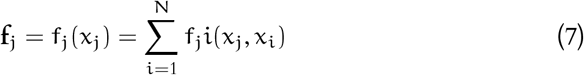
3. Given *x_j_*, |f_*j*_(*x_j_*, *x_i_*| is a monotone function of *x_i_*

The above assumptions are satisfied by almost every well-known reaction kinetics ranging from mass action, Michaelis–Menten to Hill kinetics. Further, the transfer function matrix (**G**(s)) associated with (5) and (6) can also be obtained as

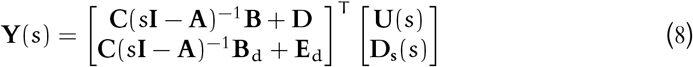

where, **D**_s_(s) is the Laplace transform of the disturbance signal **d**(t).

It is worth mentioning that the behavior of a nonlinear system around a steady-state does not always resemble its linearised counterpart. However, since adaptation is a stable response, the corresponding system matrix **A** is Hurwitz. Therefore, applying the celebrated Hartman-Grobman theorem in this case, it is possible to establish a local diffeomorphism around the operating point between the responses of the nonlinear and linearised systems [32].

### 2.2 Conditions for robust adaptation

In the language of control theory, perfect adaptation can be cast as the problem of output regulation (disturbance rejection and perfect tracking) in the presence of deterministic disturbance. Adopting a linearised representation of the nonlinear system enables us to use the wealth of regulator theory to deduce the conditions for robust, perfect adaptation.

For a linearised system in (4), (5), and (6), the conditions for perfect adaptation can be cast as a problem of designing a control input *u*(t) such that i) the closed-loop control system is asymptotically stable and ii) zero final gain 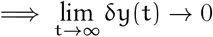. Further, the feedback control can be of two types namely i) static state feedback [33]: **u** = −**k**_*x*_**x** + **k**_*d*_**d** and ii) dynamic output feedback, which can be represented as

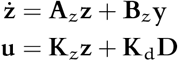

Therefore, the closed-loop system can be represented as

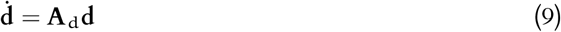

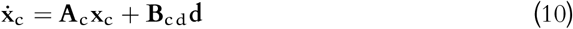

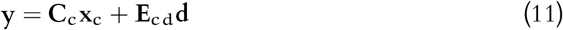

It is well-known in the literature of linear regulator theory [34] that there exists a controller u that can perform perfect regulation for the system described in (4),(5) and (6) if and only if there exists an unique matrix **P** such that the following equations are satisfied

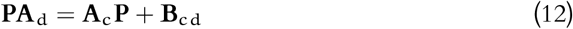

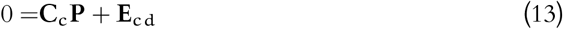

The first set or equations in (12) are also called the Sylvester equations [34]. It has already been established in the literature that there exists a solution **P** for the Sylvester equation if the matrices **A**_d_ and **A**_c_ share a disjoint spectrum. Since the control task of regulation is conditioned on stability the **A**_c_ matrix is Hurwitz. Hence, there exists at least one solution **P** for exponentially divergent or marginally bound disturbances.

Unlike the Sylvester equation, the set of equations in (13) guarantees the uniqueness of the solution. Therefore, although the existence of **P** is taken care owing to the stabilization and divergent nature of the disturbance in presence of parametric disturbance, the controller structure plays a crucial role in establishing a unique solution of **P** for the combined system of equations. In fact, it can be shown that in presence of parametric fluctuations concerning the system matrices **A**, **B**, **C** and **E**, a static state feedback can not provide perfect regulation for it renders the equations (12) and (13) unsolvable for **P** [34]. Therefore, it can be concluded that a static state feedback control strategy can not perform robust, perfect regulation. On the other hand, the *Internal Model Principle* (IMP) guarantees robust, perfect regulation with a dynamic output feedback control strategy as long as the characteristic polynomial of the controller matrix **A**_z_ is n–divisible by the same for the disturbance matrix **A**_d_ [34].

### 2.3 Towards imperfect adaptation

It is important to note at this juncture that central to the claim of the internal model principle (IMP) guaranteeing robust, perfect regulation, lies an assumption that *the controller modules (**A**_z_, **B**_z_*) are free of the parametric fluctuations. In case of a practical control system it translates to the fact that the internal dynamics of the controller and the sensor systems are not subjected to parametric fluctuations—this is clearly not the scenario for most complex networks. Therefore, we explore the prospects of *imperfect* regulation. Imperfect regulation is characterised by a non-zero finite value of the error signal—the difference between the steady-state output and the set points. In a linearized representation, perfect adaptation translates to the control problem of regulation around *zero.* Therefore, imperfect adaptation results in a non-zero, finite steady-state output. Although this is a defining feature of imperfect adaptation, it is a limiting condition, for this does not at all reflect on the control effort of regulation. For instance, any stabilizing controller, according to this condition, can be classified as an imperfect regulator. Hence, it is necessary to refine the condition for imperfect adaptation to eliminate the non-adaptive network structures.

#### Definition 1.

*A stable, LTI system with real singularities provides imperfect adaptation in presence of step-type disturbance if and only if the corresponding impulse response* (g (t)) *of the system contains at least one zero crossing in* t ∈(0, ∞).

The above definition poses a significant task before the control strategy in the presence of parametric fluctuations. Evidently, the absence of the positive impulse response throughout the asymptotic times eliminates the rise of a monotonic step response which albeit being stable, can never be a signature of regulation.

Although, the question of systems with positive impulse response (PIR) has been fairly well-discussed in the control theory literature [35–37] the relation between imperfect adaptation (regulation) and PIR has not been well-explored. Drenstig *et al* (2008) first argued that a three protein network must contain at least one zero for adaptation *i. e.* to avoid, the system being PIR [20]. Here, we propose the following sufficient condition for PIR that can further aid in constricting the search space

#### Theorem 1.

*A stable, proper, and minimum phase transfer function with real singularities exhibit PIR responses if all the zeros are situated further than the furthest pole from the origin.*

*Proof.* Given the transfer function, and the condition z_*i*_. > max(p_*k*_)∀ *i* =1(*i*)*z* and *k* =1(*i*)p. G (s) can be written as

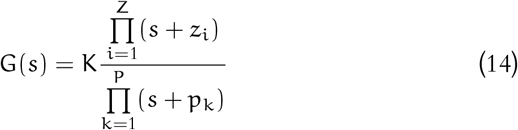

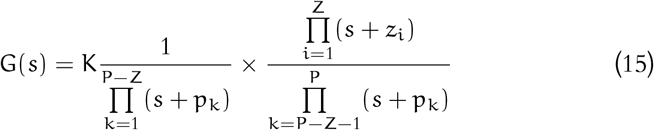

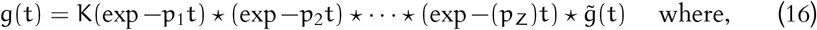

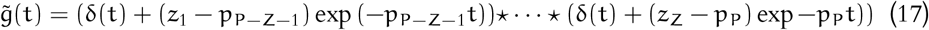

where, ‘⋆’ is the convolution operator.

As it can be seen from (16) that g(t) in this case, can be expressed as the convolution of P – Z exponential functions and 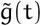. It is evident that the exponential functions are nonnegative—this renders the convolution of multiple exponential also non negative. Further, from (17) it can also be seen that all the elements of 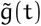 are individually positive thereby resulting in the non-negativeness of 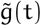. Hence, the impulse response g(t) in this case remains non-negative for all time (refer to Fig. 3).

Interestingly, it can be shown that the pole-zero position can have a direct bearing with the topological structure of the network. Therefore, casting the conditions for imperfect adaptation as a requirement of non-PIR, stable system can effectively reduce the search space in the set of structural possibilities (Supporting information).

From a design perspective, it is of great interest to synthesize systems that can mimic the response of a complex biological system. To this purpose, we present the following sufficient condition for a dynamical system to exhibit adaptation (both perfect and imperfect).

#### Theorem 2.

*A stable, causal, non-minimum, LTI dynamical system (*G(s)*) with real and simple ingularities expressed as*

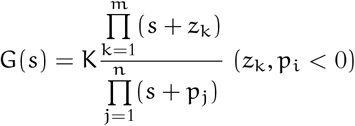

*can provide adaptation if all the zeros are closer to the origin than all the poles in such a way that 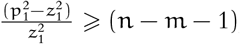, where,* z_1_ *and* p_1_ *are the closest zero and pole to the origin respectively.*

*Proof.* As per the Definition 1, both perfect and imperfect adaptation to step type disturbances require a dynamical system that involves a non-monotone step response—this implies the corresponding impulse response contains at least one finite time zero crossings. Hence, it can be concluded that the network structures capable of adaptation should contain a non-PIR (NIR) impulse response. Further, Blanchini *et al* (2017) showed that if a linear, time-invariant LTI dynamical system h(t) contains a positive impulse response then it should satisfy the following

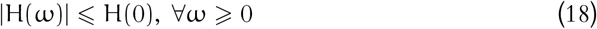

where, H(jω) is the Fourier transform of h(t) [35].

Therefore, g(t) is non-PIR (NIR) if ∃ a non-empty set Ω such that |G(jω)| ⩾ H(0), ∀ω ∈ Ω, where, G(jω) is the associated Fourier transform of g(t).

The frequency response for g (t) can be written as

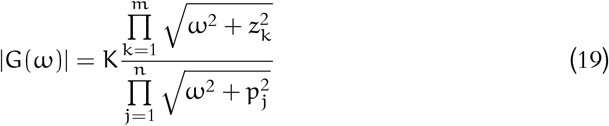

Since, |G(ω)| is non-negative, the sign of 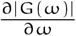 coincides with the same of 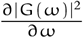 for allω.

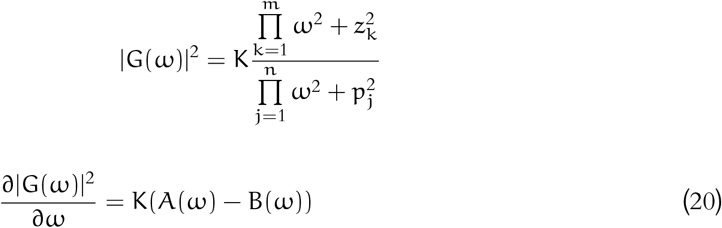

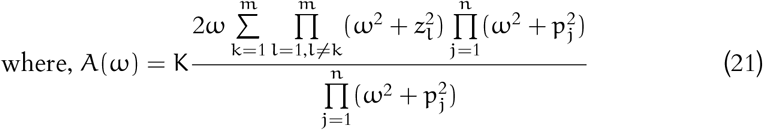

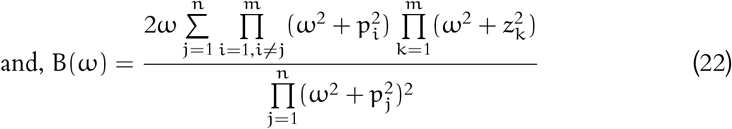

Let us define a function 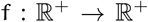 such that f(p) = ω^2^ + p^2^. Correspondingly the induced set function 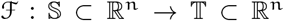 is defined as 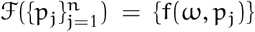. Through appropriate grouping of the elements of A(ω) and B(ω) we obtain from (21)

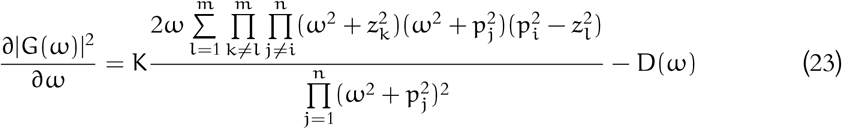

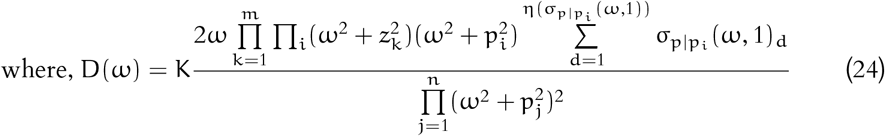

where, η is the cardinality operator, the set {i} denotes the indices of the poles that are grouped with the zeros in (23), p/p_*i*_ refers to the set of poles excluding {p_*i*_}, the set σ_p|p*i*_ (ω, 1)) contains the all possible n – η(p/p_i_) – 1 multiplicative combination of 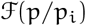.

It is evident from equation (23) that it is necessary to have at least one zero that placed before the associated pole for the initial slope to be positive. As per the assumption, the zeros are placed before all the poles. Let us consider the region 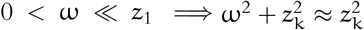 and 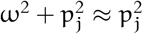.

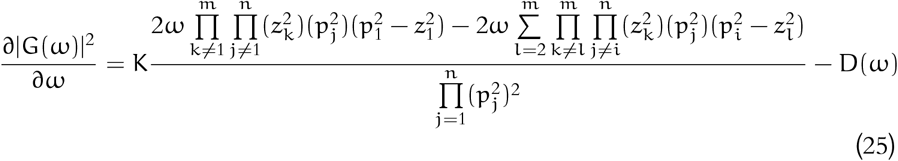

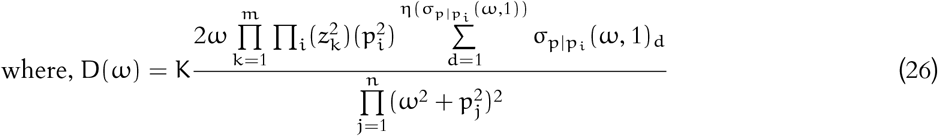

Let us define

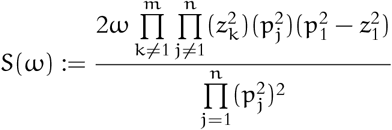

Since, z_1_ < z_2_… < z_*m*_ and p_1_ < p_2_… < p_*n*_, it is evident that the coefficient of 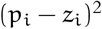 in (25) is maximum among all the coefficients of (p_*i*_ – z_*i*_)^2^. The individual signs of all these coefficients are positive (zeros are placed before the poles). Further, due to the aforementioned ordered arrangement of {z_k_} and {p_k_}, the ration between D(ω) and S(ω) for 0 < ω ≪ z_1_ can be expressed as

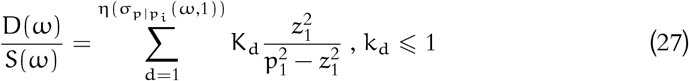

Since, it is given that 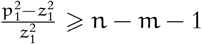 and η(σ_p|p_i__ (ω, 1)) ≤ n – 1,

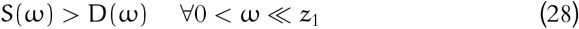

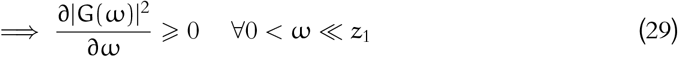

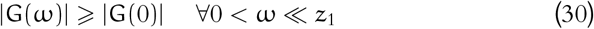

Therefore, the violation of non-PIR impulse response satisfies the condition for adaptation.

It is to be noted that the above theorem, being a sufficient condition, allows much redundancy *i. e.* the dynamical system can provide adaptation, even if the condition of all the zeros placed before the pole is relaxed. It can be shown that for a sufficiently high distance between z_1_ and p_1_, it is possible to design adaptive networks with the rest of the zeros placed after p1 (Refer to supporting information) as suggested in Fig. 3. Intuitively, the primary design idea that should be driven through this theorem is that *the likelihood of imperfect adaptation increases with increasing the distance between the zero and the pole closest to the origin.*

**Fig 3.**
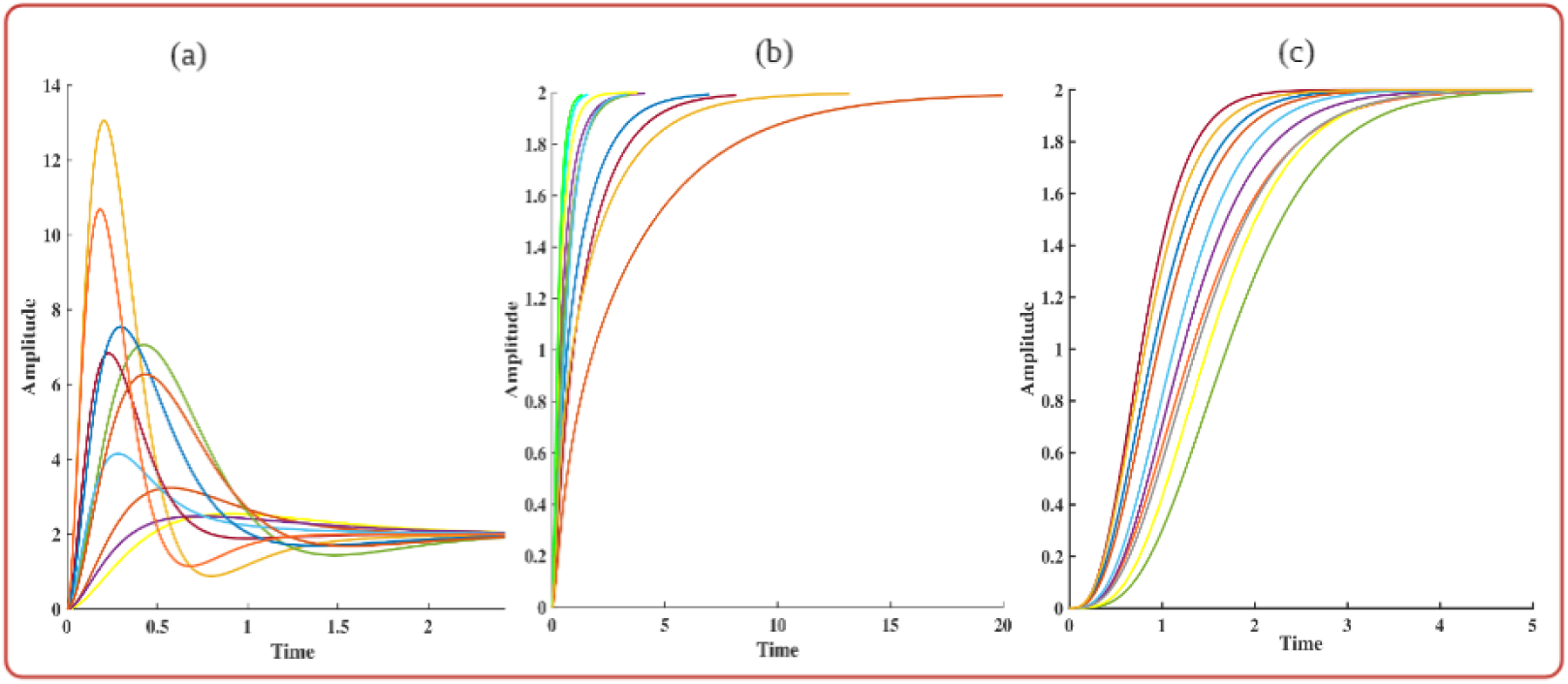
Represents the response of three classes of causal, stable and minimum-phase transfer functions where (a) at least one zero is placed before all the poles, (b) all the zeros are placed after the poles, and (c)there exists no zero. The exact pole-zero positions are chosen at random. Refer to the supporting information for the transfer function parameters.

### 2.4 Variance minimization

The notion of robustness deals with the performance of the system in the presence of parametric uncertainties—this leaves out the crucial scenario that in reality, the system at hand suffers from several disturbances that do not manifest as parametric fluctuations, for instance, unmodelled disturbances, sensor noise. Therefore, it is necessary to also observe the performance of the system in the presence of noise along with parametric fluctuations. For a dynamical system disturbed by the vector of random, incremental Brownian motion the corresponding linearized state space version can be represented as

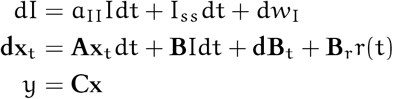

where, d*w*_I_, **dB**_t_, r(t) are incremental Brownian motion added to the network, and the reference signal respectively. The corresponding transfer function can be obtained as

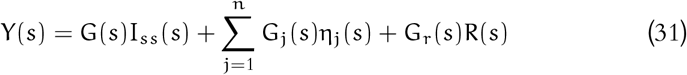

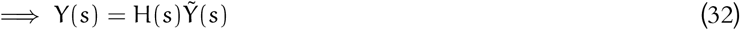

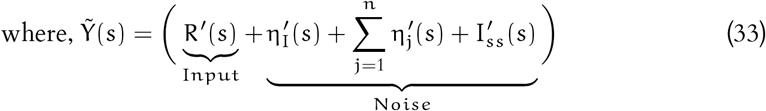

where η_j_(s) is the Laplace transform of the empirically differentiated version of the Brownian noise (**dB**_t_)_j_. It is evident from equation (33) that the output signal has a non-zero variance due to the noise contribution. Further we propose the following criterion for the well-definedness for a Wiener–Kolmogorov filter.

#### Definition 2.

*A W-K filter formulation is well-defined if the noise terms* 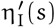 *and* 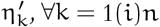 *equation contain finite variance at all time.*

With an assumption of the input signal being uncorrelated with the noise, the Wiener– Kolmogorov filter equation for minimum output variance can be expressed as [38]

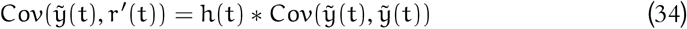

Therefore, with this equation, it is possible to design a h(t) by comparing the right and left-hand sides of the equation in such a way that y(t) follows I_ss_(t) in a minimum mean square error possible. Further, in the case where the reference signal (I_ss_(t)) is zero, then this minimisation translates to a variance minimisation program.

Further, the calculation of steady-state output variance for the system of LTI stochastic differential equations can be performed through the application of Ito’s calculus. The detailed derivation of steady-state out variance has been shown in the supporting information.

The above framework we proposed and the subsequent results aid us in building crucial insights into the structural requirements for robust adaptation in presence of parametric fluctuations and additive noise.

## 3 Applications to biochemical networks

This section presents the application of the frameworks discussed in the foregone section to biochemical networks of various sizes, ranging from two nodes to arbitrarily large networks. We first consider the scenario in which the underlying dynamics of the network is disturbed with parametric fluctuations. We use the framework of liner regulator theory to deduce the network structures for robust adaptation— this equipped us to present a systematic assessment of the performance of two commonly known adaptive topologies namely balancer and opposer modules. Additionally, we also used Theorem 2 to provide structural recommendations for improving the robustness of the adaptive modules. In the second part of this section, we deal with the exciting scenario of stochastic disturbances where we aim to minimize the steady state output variance along with the pre-assigned goal of perfect adaptation. We address the problem of variance minimization into two steps— at first, we propose a Wiener-Kolmogorov filter-based design to achieve the optimal parameters that result in minimum output variance of a given network structure. In the second step, we devise a frequency-domain based route for variance expression that enables us to compare the output variances across different network structures. The proofs for some of the propositions are kept out from the main text for the sake of brevity.

A biochemical network is constructed with the nodes representing the biochemical species (proteins, genes, metabolites *etc.*) and the edges characterising the chemical interconnection between two nodes. Further, each interaction is directed and can have two distinct impacts, namely i) Activation and ii) Repression on the target node.

In case of activation, the source species enhances the synthesis procedure of the target species, whereas repression is characterised by the act of inhibition of the same. Considering the concentration of the biochemical species as the state variables, the dynamical system underlying an N–node biochemical network can be represented as

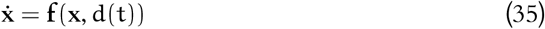

where, 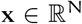 and d(t) are the states of and the external disturbances to the dynamical system, respectively.

### 3.1 Assumptions

Before proceeding to the main section of the results, we mention certain crucial assumptions on the flow of the dynamical system.

1. The set of coupled, first-order differential equations in (35) constitute a well-posed dynamical system.
2. **f** is Lipschitz with respect to the states of and disturbance to the system.
3. We also assume that the disturbance (d(t)) is mitigated through the network via single node (Let us denote it as x_1_).
4. The j^th^ node x_j_ can be written as

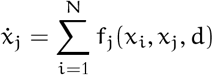
5. For given X_j_, |f_j_(d, x_i_, x_j_)| is monotonic with respect to |x_i_| or |d|.
6. The quantity 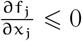 in the entire state space.

It is worth noting that because of assumption 5, the Jacobian matrix 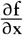 serves as digraphlike matrix for the associated network architecture. Further, the aforementioned assumptions do not point to specific rate kinetics, for most of the existing rate kinetics in the literature satisfy these assumptions.

### 3.2 Structural requirements for robust adaptation

It is well-known in the literature that perfect adaptation can be performed by two distinct classes of network structures [24, 27]: (i) negative feedback loop with buffer action or balancer modules, as termed by Araujo *et al* (2018), and (ii) multiple feed-forward paths from the input-receiving to the output node with mutually opposing effects or opposer modules as denoted by Araujo *et al* (2018). We here provide a theoretical evaluation of these network architectures vis-a-vis their performance in the presence of parametric disturbances.

#### 3.2.1 Proposed control strategy

As discussed in the methodology section, the objective of providing perfect adaptation (regulation) can only be performed by a dynamic feedback controller. Further, the output-to-controller transfer function should contain an n–copy of the disturbance model D(s). Following this, the system should contain at least one integrator to reject any step or staircase type disturbance. Therefore the dynamic error feedback control can be expressed as

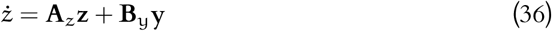

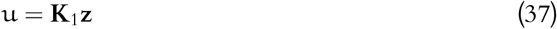

where, **z** and u are the controller state variables (often termed as auxiliary variables) and the control input respectively.

In a classical dynamic error feedback control, the equation in (36) contains the error signal (y(t) – r(t)) instead of only the output term. Since perfect adaptation translates to the invariance of the output steady-state of the nonlinear system, the reference signal in the linearised expression should be zero.

Further, the controller expression in equation (36) assumes that the controller receives information about the process directly through the output state only. This translates to the fact that the controller node has to have an edge from the output node, which is not always the case for biochemical networks. There exist ample network structures that can provide perfect adaptation without an edge from the output to the controller node. For this purpose, we propose the following control strategy that is inclusive of all possible scenarios

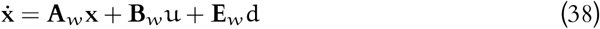

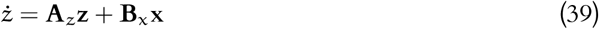

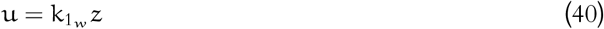

where subscript ‘*w*’ refers to the disturbed states of the matrices in the presence of parametric fluctuations. Further, we assume that the disturbance factor does not lead to the *sign change* of any elements of **A**_*w*_ and the input disturbance is mediated through the state *x*_1_.

For perfect adaptation, the proposed control strategy in equation (39) has to be brought into a form amenable to the application of the celebrated internal model principle.

##### Claim 1.

*If the pairs* (**A**_w_, **E**_w_) *and* (**A**_w_, **B**_w_) *are structurally controllable* ∀ *w then it is always possible to rewrite equation* (39) *in the Laplace domain as*

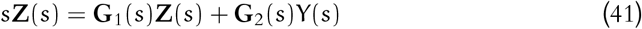

*where,* ‘s′ *is the Laplace variable.*

In a biochemical network with N—nodes, it is sufficient to consider a single node as the controller node for perfect adaptation to stair-case type disturbance (Bhattacharya *et al*). Further, as we shall see later, a controller module of a bigger size undermines the concept of robustness in biochemical networks. Therefore, in this work, we shall restrict to a single controller node.

#### 3.2.2 Structural implications

Claim 1 enables us to bring the proposed controller in equation (39) to a form amicable for applying the internal model principle (IMP). From IMP, for a step (stair-case) type disturbance, the single controller node should act as an integrator to reject the disturbance— this implies *the term* 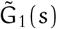 *in equation* (41) *has to be zero.* Further, from equation (4), G_1_(s) contains the term A_z_ and the contribution from the expressions of 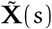 and X_1_(s). Let us decompose G_1_(s) into two parts

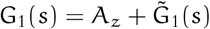

where, 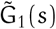 captures the indirect contribution of 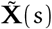 and X_1_(s) to the dynamics of the controller.

There can be only two possibilities for G_1_(s) to be zero: i) 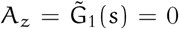 or ii) 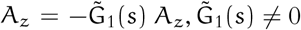

##### Proposition 1.

*The first condition,* 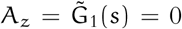 *translates to a special class of negative feedback loop with the controller node with the following properties*.

1. *The controller must perform a buffer action.*
2. *The controller node should not contain an incoming edges to any of the nodes in the loop whose concentration profile in Laplace domain can be expressed as a sole function of the output node concentration.*

##### Proposition 2.

*The second condition,* 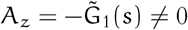 *translates to the requirement of multiple forward paths from the disturbance-receiving node to the output with mutually opposite effects.*

It is established in Proposition 1 that the negative feedback strategy with buffer nodes (balance modules) provides adaptation through a direct integral action facilitated by making A_z_ =0 and further specific structural adjustments as shown in Fig. 4. Therefore, if the controller parameter A_z_, is not vulnerable to any disturbances, then *negative feedback loop exhibits perfect adaptation in the presence of the disturbances in the process parameters*.

**Fig 4.**
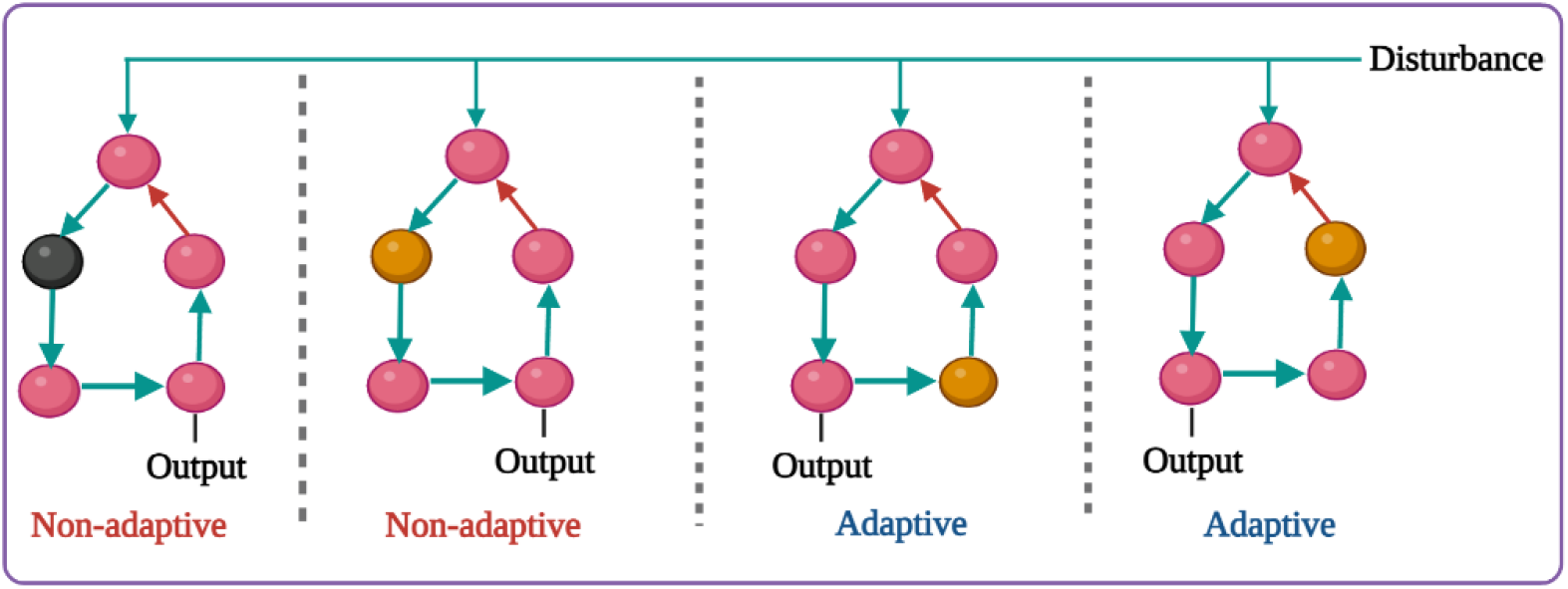
Structural implications of Proposition 1. The black node in the left most network refer to the controller node without a buffer action rendering the feedback loop incapable of adaptation. On the other hand, the nodes with brown colour refersto buffer action. The nodes in Pink refer to the process modules. Further, as per proposition 1, not all negative feedback loops with buffer action can produce perfect adaptation.

On the other hand, from Proposition 2, it is clear that the multiple feed-forward structures with mutually opposing effects (opposer modules), as shown in Fig. 5, require 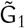—the contribution from the process to the integral action—to be non-zero. Therefore, the fluctuations in the parameters pertaining to the process module also seep into the controller via 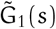 rendering the pure integral action an impossibility. Hence, *mutually opposing feed-forward structures can not provide perfect adaptation in the presence of parametric fluctuations in the process modules*

**Fig 5.**
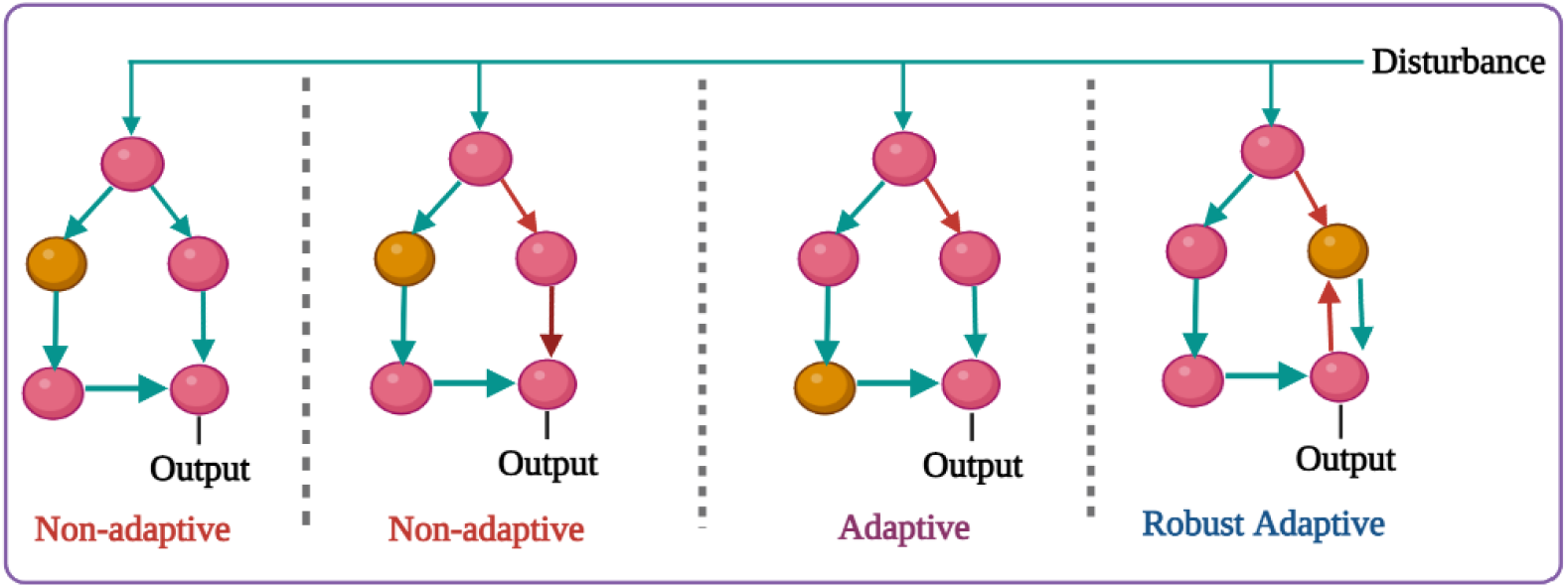
Structural implications for Proposition 2 and 3. The brown node in the left most network refer to the controller node. The nodes in Pink refer to the process modules. Further, as per proposition 3, addition of a negative feedback improves the robustness of IFFLP in presence of fluctuations.

**Fig 6.**
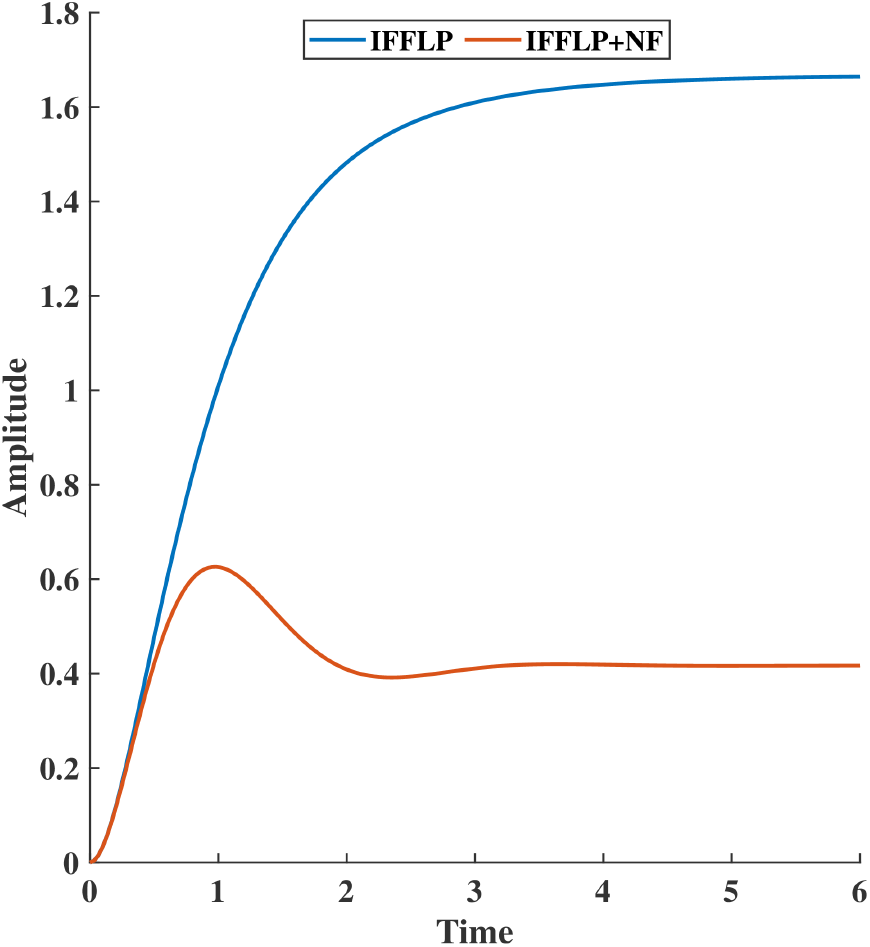
Addition of negative feedback improves the performance of IFFLP. The system parameters necessary for the simulations are provided in the supporting information

#### 3.2.3 Improving the performance

The opposer modules distribute the task of accomplishing perfect adaptation between both the process and controller modules which makes it vulnerable to random parameter fluctuations—this can also serve as a theoretical justification for the scarcity of opposer modules as adaptation-capable topologies in nature. Therefore, it becomes essential for synthetic design to provide structural recommendations to better the performance of the opposer modules in the presence of parametric fluctuations.

##### Proposition 3.

*The addition of a negative feedback loop to a stable opposer module pushes the pole closest to origin towards* – ∞.

Although it can be tempting to appreciate Proposition 3 in the light of the well-known result on the use of feed-forward and the output feedback strategy for improving the relative stability the fact that Proposition 3 does not pose any condition on the position of the feedback loop. For instance, even if the negative feedback loop is constructed through the nodes pertaining solely to the process module (not the controller) the statement for Proposition 3 stands true.

##### Remark 1.

*As established in Proposition 3, addition of negative feedback pushes the pole nearest to the origin further left. We have also argued in the foregone sections that placing the zeros before the poles can potentially serve as a sufficient condition for adaptation. It is quite well-known that perfect adaptation requires the zeros of the stable system to be placed precisely at the origin. Therefore, in the presence of parametric disturbance, when the zero position is deviated towards left from the origin, the addition of negative feedback to the opposer node provides crucial support for it increases the distance from the zero and its nearest pole thereby retaining the adaptation-capability of the motif (refer to Fig. 6). Further, the zero positions remain unchanged if the negative feedback does not introduce any new forward paths from the disturbance-receiving node to the output node.*

Intuitively, the conclusion of Proposition 3 is also applicable for balancer modules. It can be shown that the net increment in the determinant of the closed-loop system matrix is positive as long as the additional P—node negative feedback shares at least one common edge or node with all the existing loops. Further, all the coefficients of the s^N–*j*^∀, j = p(*i*)N in the characteristics polynomial of the closed-loop system matrix increases post addition of the negative feedback.

### 3.3 Adaptation in presence of noise

We hitherto assumed a deterministic setting to deduce the design principles for adaptation. However, apart from the inherently stochastic nature of biological networks, the classical approach toward finding network structures for particular functionality has always assumed that the network under consideration is dedicated exclusively to performing the functionality. Although this abstraction aids in drawing great insights into the relationship between network structures and specific functionalities, it neglects the unmodelled connection patterns. To circumvent this problem, we assume that each node has a source of randomness from unknown connection patterns. The resulting dynamical system can be represented as

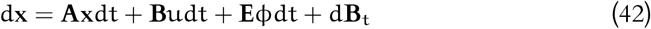

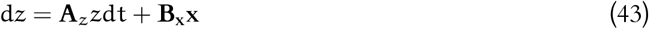

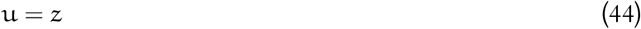

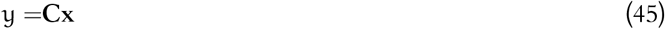

where, ϕ(t) is the external disturbance.

We propose a two-level methodology to explore the problem of’ *variance minimization.’* Firstly, we set out to answer the following question: how does an adaptation-capable motif perform in terms of the steady state output variance compared to its closest neighbor motif across the parameter sets? In the second step, we propose a W-K filter-based mechanism to minimize the variance for a given network structure.

#### 3.3.1 Variance minimization across the structure

It is evident that the output variance of a network is dependent on both the structure (interconnections) and the associated parameters (rate kinetics) of the associated dynamical system. In the presence of parametric fluctuations, structure plays a crucial role in minimizing the variance across different topologies. In this section, we compare the steadystate output variance for each class of the network structures (balancer and opposer modules) with their respective nearest neighboring structures.

##### Definition 3.

*Nearest neighboring network structure: Two network structures 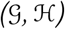 are called nearest neighboring network structure if the associated digraph matrices of 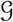 and 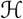 only differs by the sign of a single element.*

According to the above definition, the nearest neighbor of a balancer module can contain a positive feedback loop (involving the same nodes) along with the buffer action or a negative feedback loop without the buffer action. It is to note here that the buffer action is facilitated by making a diagonal component associated with the (linearised) dynamics of the controller node zero.

Further, due to the instability of the positive feedback loops with buffer node [27], we start with the network structures containing negative feedback loop without buffer node. We first start with a 2–node balancer module and subsequently, extend the results to the N–node case.

##### Claim 2.

*A* 2–*node balancer module 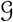 provides maximum variance across all the network structures that 1) belong to the set nearest neighboring set of 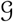 and 2) results in a Hurwitz digraph matrix.*

*Proof.* In case of a 2–node balancer module, with the digraph matrix **A** = [a_11_ a_12_//a_21_ 0], 0, the transfer function relation between the adaptive output and all the disturbances can be written as

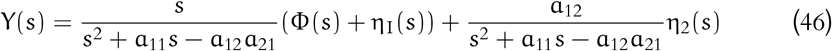

Further, the power spectral density of the input and the output is related in the following manner

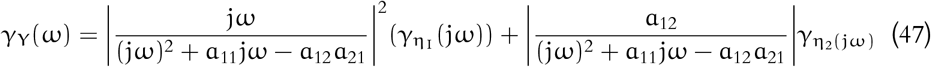

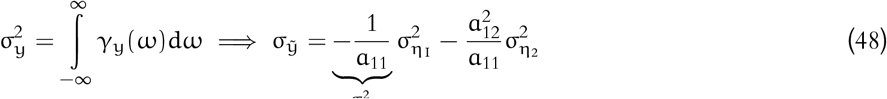

where, the above equations assume the uncorrelatedness amongst the noise signals introduced at different nodes and the minimum-phase transfer functions.

For the non-buffer scenario a_22_ < 0, the output variance can be obtained as

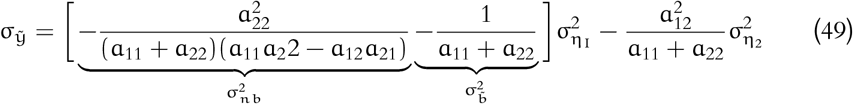

Since, a_ii_s, ∀ *i* = 1(*i*2) are negative and a_21_a_12_ < 0

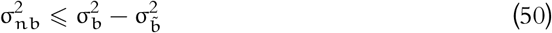

Further, due to the simple fact that –(a_11_ + a_22_) > –a_11_ the contribution of η_2_ to the output variance is also reduced. Therefore the 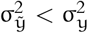. This concludes the proof.

##### Remark 2.

*Although the above Claim works with* 2—*node balancer modules. The same result can be generalized to the* N—*node scenario as well. The balancer module, as established in the foregoing sections, attains perfect adaptation through the buffering action* (a_jj_ = 0*, where* a_jj_ *is the diagonal element corresponding to the controller node* **x_j_**) *i. e. the buffering action places the closed loop zero on the origin. Therefore, violation of buffering action* (a_jj_ < 0) *result in a zero placed at* a_jj_ *instead of the origin. Further, addition of a negative diagonal element on the closed loop system matrix also increases all the coefficients in the characteristics polynomial of the system matrix **A**. To illustrate the fact, we consider two* N^th^ *order transfer functions* G(s) *and* H(s) *related in the following way*

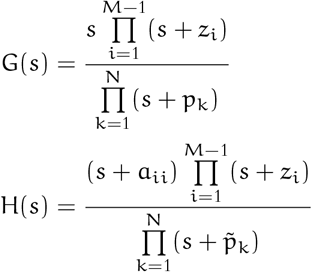

*The output signals associated with* G(s) *and* H(s) *are* Y(s) *and* 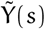 *respectively. Given the input variance to be* 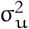 *the associated variance expression can be obtained as*

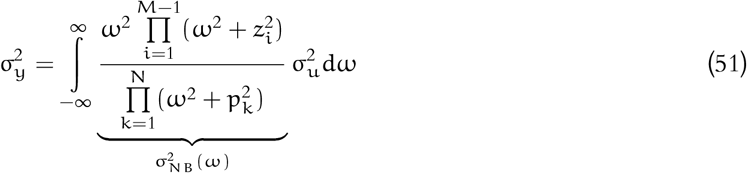

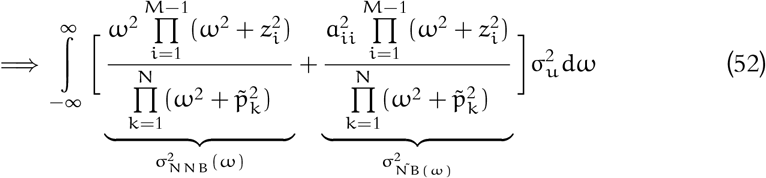

*Using combinatorial matrix theory it can be shown that*

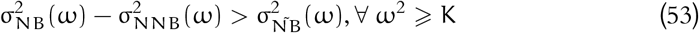

*where* K *is a finite constant. Since, the variance integrates between the entire region i. e.* ω ∈ (–∞, ∞) *the following holds true*

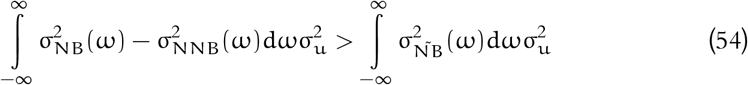

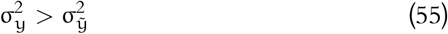

*Therefore, an* N–*node,* N–*edge balancer module attains perfect adaptation at the cost of variance higher than all its stable neighboring network structures.*

The opposer modules, on the other hand are shown to provide perfect adaptation through the multiples mutually opposing forward paths from the input disturbance-receiving node to the output. Therefore, the nearest neighboring network structure (that produces a Hurwitz digraph matrix) of an opposer module is the class of feed-forward structures containing multiple forward paths with same (coherent) effect on the output node. The following claim compares the steady-state variance of the output node for an opposer module with that of its neighboring topologies.

##### Claim 3.

*An* N–*node*, N–*edge opposer module, fully controllable by the disturbance input provides minimum variance across all its nearest neighboring network structures that yield a stable, non-minimum phase system.*

*Proof.* An N–node feed-forward module contains a lower triangular digraph matrix. Therefore, the associated system poles are the diagonal elements. Since, none of the stable nearest neighboring motifs tinker with the diagonal elements the dynamical systems associated with the opposer module and its nearest neighboring topologies share identical eigenvalues. As a result of this, the poles of the transfer functions remain unchanged if one traverses from the opposer module to its nearest neighbors– a set of feed forward structures with all coherent forward paths. However, the zero positions owing to a certain class (denote as 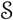) of transfer functions changes owing to the introduction of coherent forward paths in the network. The set 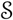 denotes all the nodes that has multiple forward paths to the output node. Unlike the opposer modules, the feed forward network with coherent forward paths shall shift all the zeros pertaining to the transfer functions between the output and the noise added to the nodes in S from the origin towards the left half of the s—plane. The final output expression for the opposer module (Y(s)) can be obtained as

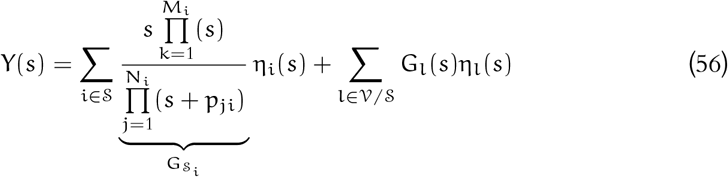

where, the 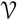 is the set of all vertices. For all nodes in the set 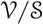 the transfer functions 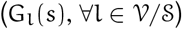 shall remain across different feed forward network structures including the opposer modules. Therefore, the output 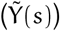 for the neighboring structures can be expressed as

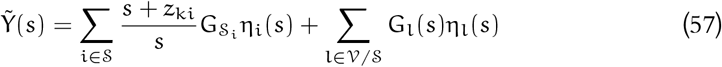

Comparing the variance of the output for the opposer module 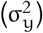 with that of the output from the coherent feed forward structures 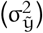

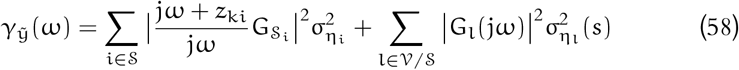

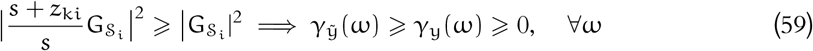

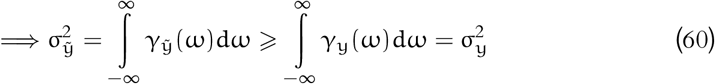

Therefore, as the above equation suggests, the opposer module structurally minimizes the variance across its neighboring network structures.

Fig. 7 provides an interesting computational study that validates the theoretical findings in Claim 2, Claim 3, and Remark 2. In the case of the balancer node the buffering action is violated through making the buffer diagonal non-zero negative. The variance is calculated in each scenario. As suggested by Claim 2 and Remark 2, the variance decreases monotonically as the buffering action gets violated. Similarly, as suggested by Claim 3 the variance remains minimum for the opposer module.

**Fig 7.**
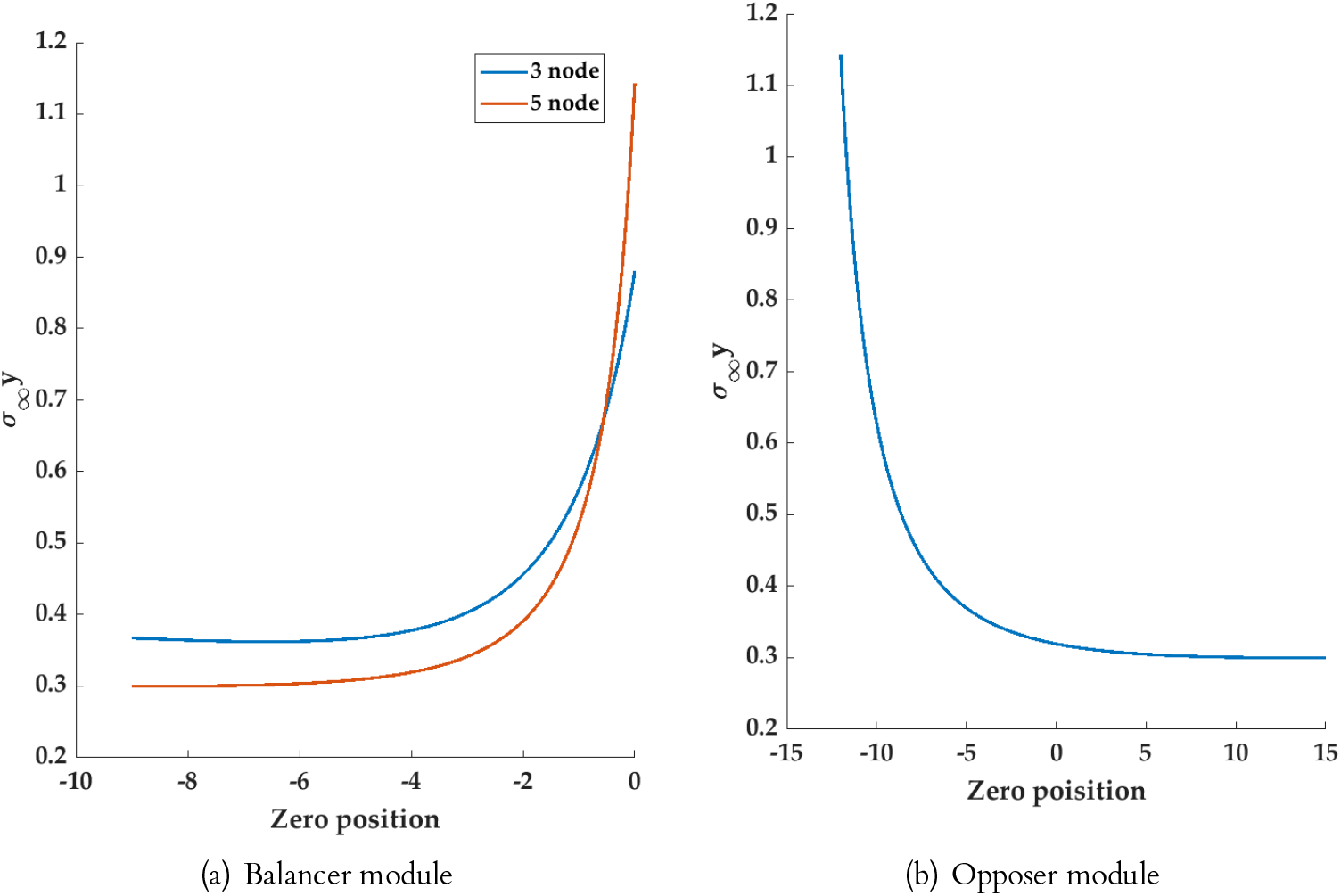
Plot of steady-state output variance concerning the zero position. (a) showcases the increasing trend of the variance as the network tends towards the balancer module. (b) shows the capability of an opposer module to minimize the steady-state output variance across its neighboring network structures. The positive zero position refers to a non-minimum phase system that can be ruled out by design. The theoretical method for calculating the variance has been detailed in the supporting information

#### 3.3.2 Adaptive motifs as Wiener-Kolmogorov filter

The previous section enables us to compare the steady-state output variance across the network structures whereas in this section, we explore the domain of the parameters while fixing the network structure. For this purpose, we adopt the celebrated W-K filtering algorithm to fine tune the parameters. At the same time, as per Definition 1, application W-K filter filter requires a feasibility check of the network structures. Therefore, we examine whether the given network structures of interest *i. e.* the balancer and opposer modules run into any feasibility issue in serving as candidate W-K filters.

Consider a two-protein network where two proteins 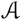 and 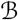 are connected with each other. The concentration of 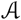 and 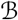 are denoted as x_1_(t) and x_2_(t). Further, since **A** is the output node, it is safe to assume that the node 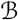 serves the purpose of the controller node (integral action). The dynamics can be written as

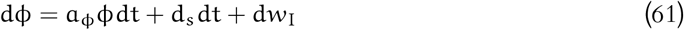

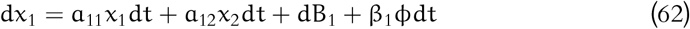

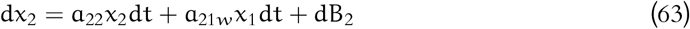

As it is evident, we first assume that there exists no parametric disturbance. The consideration of fluctuations in parameters shall be incorporated in the final analysis.

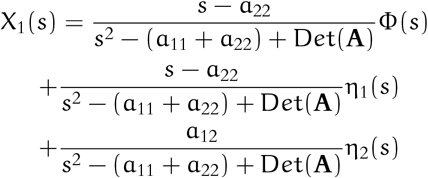

where, Φ(s) is the Laplace counterpart of ϕ(t) and η_1_(s) and η_2_(s) the Laplace transform of the empirically differentiated version of dB_1_ and dB_2_ respectively. Taking the highest common factor across the denominators of the transfer functions associated with η_1_(s) and η_2_(s) the output can be expresses in a W-K compatible form as

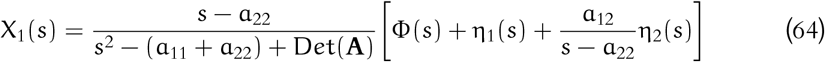

Further, it can be easily shown that for the steady-state mean of the output state to be zero, the term a_22_ has to be zero. Therefore, the output expression in the scenario of perfect adaptation

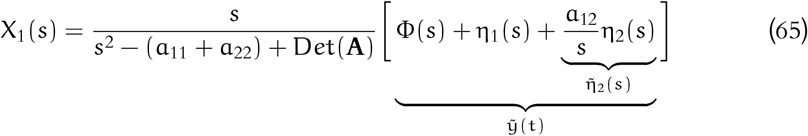

As it is evident from equation (65), for a constant disturbance d_s_, 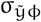 is always a exponentially decaying quantity (Supporting information). Therefore, from the W-K filter equation

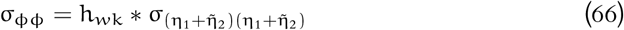

For a stable, causal, infinite impulse response filter, with distinct singularities h_*w*k_(t) can always be expressed as weighted sum of N exponentials

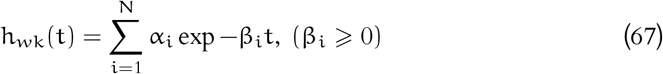

Further the quantity 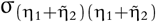 can be written as

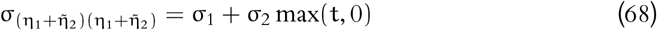

Comparing the above equation with equation (65), the right-hand side of equation (65) contains an exponentially decaying term, whereas the left-hand side contains the convolution with exponentially decaying terms with the term σ_2_ max(t, 0) which is *diverging.*Therefore, it can be concluded that there exists no feasible solution for a 2^nd^ order filter for equation (65) in the given structural setting.

##### Claim 4.

*An* N–*node balancer module capable of perfect adaptation can not serve as an* N*^th^ order* (N *poles) W-K filter irrespective of the position of the output nodes.*

*Proof.* In the case of N–node, N–edge networks, as established earlier, the balancer module involves at least one loop to attain perfect adaptation. Therefore, the multi-input,multi-output transfer function for the output Y(s) and system inputs I(s), R(s) and η_i_(s), ∀i = 1(*i*)N to share a common denominator-the characteristics polynomial of **A**. Further, in the case of balancer modules, the output node (k^th^ node) preserves the quality of perfect adaptation (in the sense of expectation) even if the disturbance is applied on the output node. Therefore, the transfer function between the output (Y(s)) and the noise η_k_(s) applied on the output node contains an (N – 1)^th^ order numerical polynomial with a singularity at the origin. The resultant transfer relation can be written as

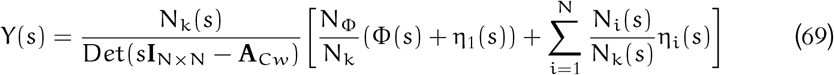

As it can be seen, since the degree of N_k_(s) is the maximum across the numerators, the resultant Wiener–Kolmogorov equation becomes infeasible due to divergence issue owing to the singularity of N_k_(s) at the origin. This concludes the proof.

Although the balancer modules can not be treated as N^th^ order filters, it is possible to work with a reduced order. For instance, instead of conceiving an N^th^ order filter, it is possible to construct a filter by only considering a factor of the term Det(s**I** – **A**) in Eq. (69). Further, the filter coefficients can be obtained through Eq. (66).

The opposer modules, as discussed earlier, provide perfect adaptation through opposing feed-forward paths from the disturbance-receiving node to the output node. Due to the absence of any loop, it is possible to write the associated system matrix **A**_Cw_ in a lower triangular form, assuming all the nodes are controllable by the input-receiving node—this inspires the following Proposition.

##### Claim 5.

*It is always possible to design an* N–*node opposer module capable of perfect adaptation as a fine-tuned first-order W-K filter.*

*Proof.* Let us assume the output is measured as the concentration of the N^th^ node without any loss of generality. For an opposer module with the network being full-structurally controllable by the disturbance input ϕ(t), ϕ(t) by being connected to only x_1_(t), and the controller node being the k^th^ node, the MISO transfer matrix for the output can be expressed as

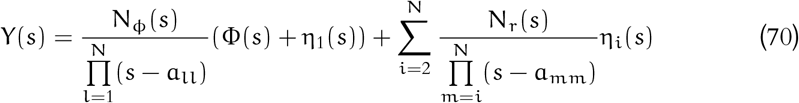

where, a_ii_ is the diagonal element associated with the i^th^ row of of **A**_Cw_.

Therefore, the highest common factor across (HCF) the denominator of all the transfer functions in the equation (70) is (s +a_NN_). Further, in all the transfer functions with zeros, the difference between the degree of the denominator and numerator polynomials is greater than or equal to two. Therefore, taking the HCF of the denominators out of every transfer function retains the strictly proper characteristics.

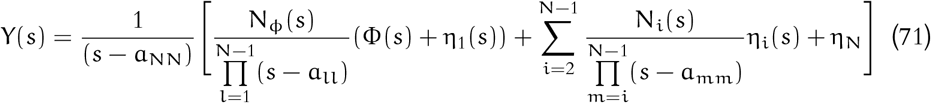

Therefore, it is evident from the above equation that the filter does not run into any feasibility issues. This concludes the proof.

It seems from the above analysis that the adaptive module enables us to avoid the feasibility issues at the cost of reduced degrees of freedom. Therefore, even in the case of the balancer modules which have a tendency of providing higher variance (from Claim 2 and Remark 2), the parameters of the balancer modules can be fine tuned using the W-K filter formalism to reduce the variance.

## 4 Discussion

The inherent diversity of biochemical networks leads to several emergent phenomena crucial to every living organism’s survival. Therefore, unraveling the structural underpinnings for the governing dynamics of the biological system is instrumental toward an understanding of biological systems. Subsequently, understanding the structural mechanisms for achieving robustness is not only central to the problem of synthetic design but also to formulating a scientific perspective about the relationship between evolvability and robustness.

It has been well-established in the literature that only the balancer and opposer modules can provide perfect adaptation. Recently, Bhattacharya *et al* (2022) also argued that the balancer module should contain at least one negative feedback engaging the buffer node to provide perfect adaptation [27]. Despite certain seminal contributions in this domain, the question of adaptation in the practical scenario *i. e.* imperfect adaptation has remained relatively unexplored. Although Ma *et al* (2009) and Otero-Muras and Banga *et al* (2019) did perform a computational screening approach, a proper theoretical intervention for the generic results has been awaiting [4, 16]. In continuation with the same, a theoretical evaluation of the admissible network topologies for adaptation also required scholarly contributions from the intersections of control theory and systems biology.

Further, it should also be kept into account that the entire exercise of discovering the design principles for a given functionality is based on a critical assumption—the given biochemical network is only dedicated to performing the particular functionality in consideration, which is far from reality. The suggested motifs for a given functionality are connected to a big network that may also facilitate cross-talk across different cells as well. At the same point of time, *no model is perfect* because, in full extension, a perfect model for a biochemical network requires modelling the entire living organism, which is not at all feasible for the given task. With the ever-increasing size of the network, it becomes a daunting task to implement a computationally involving approach to draw insights into the relationship between the structure and the governing dynamics. Therefore, an elegant, theoretically sound abstraction is required that not only aids in theoretical investigations but also guides the computational study toward a realistic scenario.

The present paper addresses the first challenge of imperfect adaptation from the systems-theoretic approach. We, for the first time, argued that adaptation requires at least one zero-crossings in the impulse response of the underlying dynamical system—this aids in connecting the design requirements for imperfect adaptation to the studies on everywhere positive (negative) impulse response (PIR) systems. The current results in the study of PIR systems resulted in a significant reduction in the search space of network structures that was achieved by Drenstig *et al* (2008) [20]. Further, we proposed another sufficient condition in Theorem 1 for PIR that translates to a further round of elimination of network structures in the search for adaptation-capable topologies. Additionally, for the first time, we proposed a sufficient condition via Theorem 2 for systems that can produce imperfect adaptation—this was crucial in our quest to provide structural suggestions for improving robustness.

We used a linear regulator theory approach to unravel the design principles for robust, perfect adaptation. To this extent, we proposed a dynamic state feedback controller that caters to the diversity of the biochemical network structures. Claim 1 establishes the compatibility of the proposed control scheme vis-a-vis the internal model principle, which is central to achieving perfect adaptation. Further, Proposition 1 suggests that the well-known balancer module performs perfect adaptation by assigning the role of disturbance rejection solely to the controller node. As long as there exist no fluctuations in the parameters concerning the controller module, the balancer module provides perfect adaptation. Therefore, *Balancer modules are robust to parametric fluctuations in the process nodes.* On the other hand, Proposition 2 points to the curious case of opposer modules that attain perfect adaptation by distributing the task of disturbance rejection to both the process and controller module. In this sense, the opposer modules are not robust to the fluctuations of either the process or the controller parameters.

Although the balancer node is touted to be the most robust network structure with respect to parametric fluctuations, it assumes the parameters describing the dynamics of the controller node to be free from any fluctuations. There is no reason to assume a controller with undisturbed parameters, for, in reality, the controller is also a node in the same biochemical network. Therefore, it is safe to conclude that *no adaptation-capable network motifs can provide robust, perfect adaptation with respect to overall parametric fluctuations*. This motivated us to investigate the prospects of imperfect adaptation. Together Proposition 3 and Remark 1 establish the fact that the addition of a negative feedback loop can increase the extent to which the adaptation-capable modules provide adaptation (perfect or imperfect) in the presence of overall parametric disturbance.

Further, we provided a theoretical abstraction to deal with the inaccuracies due to the unmodelled connections. We add an incremental Brownian noise to every node to encounter the missed connections. In this scenario, our objective becomes twofold i) deduce network structures that can provide perfect adaptation in the sense of mean—this is equivalent to finding adaptation-capable structures in the deterministic scenario and ii) reduce the effects of the unmodelled connections to its adaptive response. The second objective we approach the second objective in two subsequent steps. First, we addressed the task of exploring the structural aspect of the variance minimization problem, we proposed Claim 2 and Remark 2 that shows the relative incapability of the balancer modules to act as a variance minimization motif while retaining perfect adaptation-This can serve as the primary reason behind the poor performance of a pure antithetic integral controller– a class of balancer modules proposed by Khammash *et al* (2016,2018) [17, 18]. On the other hand, as the Claim 3 suggests, the opposer modules provide a comparative structural advantage for it minimizes the variance across its neighboring structures. Moreover, although the balancer modules are more robust to parametric disturbances the opposer modules can potentially show case an improved performance in the presence of noise. Additionally, introduction of a negative feedback to a balancer module can partially compensate for the increased variance due to the net increment in the absolute value of the denominator of the transfer function. In the Second stage, we used the Wiener-Kolmogorov filter theory to minimize the output variance subject to the parameters of a given network structure. In this aspect, Claim 4 and 5 resolve the feasibility issue concerning the W-K filter and guarantees a feasible solution to the filter design problem.

Last but not least, the structural predictions obtained in the present work are valid for almost every form of rate reaction of the network. Therefore, the structural recommendations are generalizable to a diverse set of biochemical networks, which aligns with the hypothesis concerning the conservation of design principles. Additionally, the obtained results are not restricted to networks of a particular size, thereby providing the essential property of scalability. The methodology works with a linearized framework which restricts the results obtained to the case when the disturbance is of small magnitude *i.e.* as long as there exists a diffeomorphism between the nonlinear system and its linear counterpart and secondly, there exists no bifurcation region in the nonlinear system with respect to the disturbance variable. Therefore, a careful investigation of the actual nonlinear dynamics to unravel the structures for robust adaptation in the presence of noise and disturbance with arbitrary amplitude remains an intriguing area of future study.

## Acknowledgements

PB acknowledges funding from the Ministry of Human Resources, India.

## Supporting Information

### 0.1 Proof of Claim 1

:

*Proof.* Since the external disturbance d(t) has a direct effect on the dynamics of only x_1_ equation (38) can be rewritten as

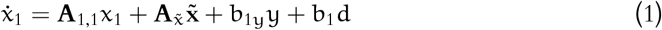

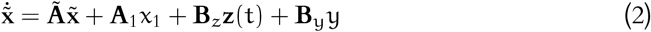

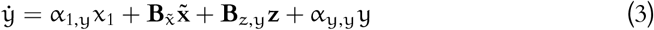

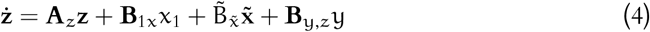

where, 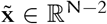 are the states apart from the input-receiving state and the output state (y). Transforming the set of equations (2)–(4) in Laplace domain we obtain

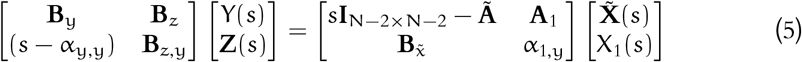

Since, the disturbance d is only applied to the node *x*_1_, the controllability condition for the pair (**A**_*w*_, **E**_*w*_) translates to the same with the pair 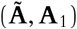. Therefore the matrix 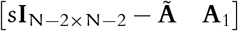 is always structurally full rank (P-B-H condition for controllability). The only way the system of linear equations in equation (5) shall have non-unique or no solution if the last row of the co-efficient matrix of 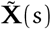 and X_1_(s) is zero which is in contradiction with the output controllability assumption. Therefore, in case of full controllability, 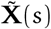 and X_1_(s) can always be expressed in terms of 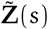 and Y(s). Further, using equation (4), it is always possible to write Z(s) in the desired form

#### Remark 3.

*The strict assumption of full structural controllability of the pair* (**A**_w_, **E**_w_) *can be relaxed further. Let us assume that there exists a module including* K *interconnected nodes that is not controllable by node* x_1_ *but is influenced by the control module **z**. Therefore, the corresponding Laplace equations can be written as*

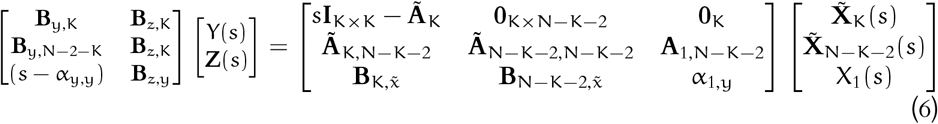

*From the* equation (6), *it can be seen that* 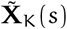 *can be solved from the first* K *equations. Therefore, the solution* 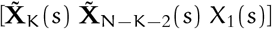 *is unique if and only if there exists at least one element in* 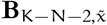 *or* α_31_ *is non-zero—this translates the the claim 1 can be made possible even when the network is not fully controllable by the external disturbance* d *given the output is controllable by the same and the ‘uncontrollable-by-disturbance’ modules are controllable by the proposed control input.*

### 0.2 Proof for Proposition 1

*Proof.* In the first case, A_z_ = 0, therefore the system matrix for the augmented state space **x**_c_:= [**x** z]′ in the closed-loop system

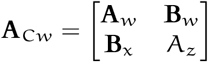

Since perfect adaptation implies stability, the matrix **A**_C*w*_ has to be Hurwitz. This translates to a set necessary conditions that require all the coefficients of s^k^, ∀k =0(*i*)N – 1 of the monic characteristics polynomial P_A_Cw__ (s):= Det(s**I**_N×N_ – **A**) to be positive. Therefore, the determinant condition for stability can be written as

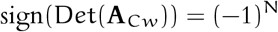

From combinatorial matrix theory, each of the N! terms in the determinant expression of **A**_C*w*_ contains a multiplicative combination of either the diagonals or the combination of off-diagonal elements in such a way that these off-diagonal combinations map to a loop (cycle) in the actual network structure [39]. Therefore, in the case where one of the diagonal elements (A_z_) of the closed-loop system matrix **A**_C*w*_ is zero, all the terms in the determinant expression of **A**_C*w*_ contain at least one loop. Further, Bhattacharya *et al* (2022) proved that in the case of buffer action (one or more diagonal elements of **A**_C*w*_ is/are zero), the network has to contain at least one negative cycle to achieve stability. This concludes the first part of the proof [27].

Further, as the condition suggests, only the negative feedback with buffer action (*i. e.*A_z_ = 0) is not sufficient for the mentioned network structure and does not guarantee a zero 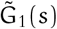 that is crucial for producing an integral control action. 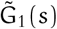 can be made zero if the (concentration of the) set of process nodes having a direct edge to the controller (denote as 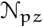) can be expressed as a sole function of the output node (Y(s)). This, in turn, is possible in an P—node, P—edge network, if there exists no incoming edge from the controller either to any of the paths from the process nodes in 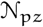 to the output node or from the output to the process nodes in 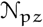. Fig. 4 illustrates the claim.

### 0.3 Proof of Proposition 2

*Proof.* For the closed-loop system, the transfer function between the output (Y(s)) and the disturbance signal (D(s)) can be written as

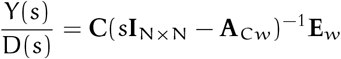

Let us denote the concentration of the K^th^ node as the output variable. Because the **E**_w_is a unit vector along the direction of *x*_1_, it can be shown that the numerator polynomial of the transfer function can be derived just by computing the minor of the (1, k) element of (s**I**_N×N_ – **A**_C*w*_)^-1^. The denominator is the characteristics polynomial of **A**_C*w*_ in terms of the Laplace variable s. Similarly, the transfer function between the controller node (Z(s)) (denote as the j^th^ node) and the disturbance input

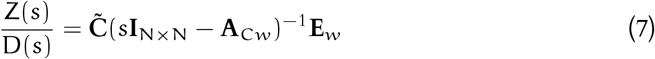

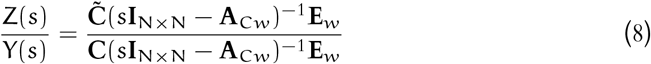

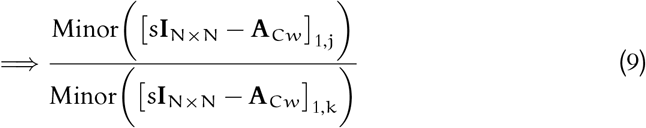

It is worth noting that the coefficient of s^0^ in the denominator coefficient is the minor of [A_Cw_]_1,k_. For the controller to provide the integral action, the denominator polynomial should be divisible by s—this translates to the requirement of the M = Minor ([**A**_C*w*_]_1,k_) to be zero. Further, the quantity [**A**_C*w*_]_1,k_ × M is an element in the determinant expression of **A**. Therefore, the quantity M encodes all possible forward paths, multiplied by the diagonal elements, from the node x_1_ to the output node x_k_. Now, in the first case, when A_z_ = 0 all the elements of M containing the diagonal element A_z_ go to zero. Therefore, through a further choice of edges, it becomes feasible to make all the elements in M individually zero without making the system structurally uncontrollable. Unlike in the first case, if A_*z*_ ≠ 0, there exist multiple elements in M. Following this, two forward paths from the input-receiving to the output nodes with opposing effects (the diagonals are assumed to be non-positive). This concludes the proof. Figure 5 illustrates the claim.

### 0.4 Proof for Proposition 3

*Proof.* An opposer module, as defined in the previous sections, does not contain any cycle—this implies that the eigenvalues of the system matrix associated with the opposer modules are the diagonal elements. Therefore, the corresponding characteristics equation of **A**_C*w*_ in case of an opposer module can be

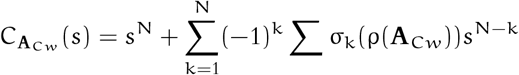

where, σ_k_ is a k-length permutation operator, σ_k_(ρ(**A**)) returns the set containing all possible k–length multiplicative combinations of diagonals from the spectrum of **A**_C*w*_(ρ(**A**_C*w*_)). Evidently, Σ σ_k_(ρ(**A**_C*w*_))s^N–k^ refers the sum across the the set σ_k_.

After addition of a negative feedback loop involving P nodes, the corresponding characteristics polynomial of the modified closed-loop matrix 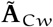 can be written as

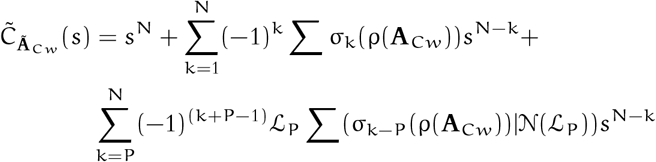

where, 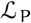 encodes the P-node loops, 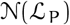 returns the nodes (rows of 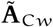) involved in 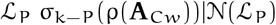 returns the set of k – P length multiplicative combination of diagonals of the rows other than the ones occupied by the loop. Since sign(L)_P_ = −1, the term 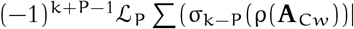 is always positive. Without any loss of generality, we assume that the diagonal a_NN_ is of the smallest magnitude among the spectrum of **A**_C*w*_ (Note, all the diagonals are by design negative). Therefore, using the concept of relative stability we can draw valuable insights about the change in the the pole positions after addition of the loop. Replacing s = s′ + a_NN_ in the modified characteristic equation

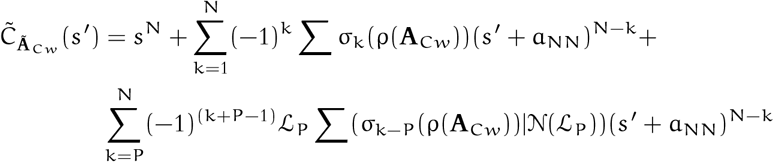

Further, through appropriate grouping of the terms (Supporting information) 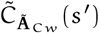 can be expressed as

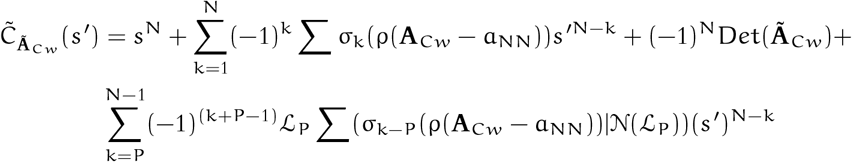

where ρ(**A**_C*w*_ – a_NN_) returns the set of all the diagonals of **A**_C*w*_ (except a_NN_) deducted by an amount of a_NN_. Since, |a_NN_| is the minimum across the diagonals of 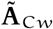 the quantity an a_*ii*_ – a_NN_ remains negative where a_*ii*_ is any other diagonal of 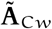. Further, because of the negative feedback the coefficient of s^′0^ remains positive. In fact, it can be shown that 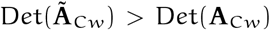. Therefore, since all the coefficients of 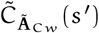 are positive, it can be shown that the magnitude of the closest root of 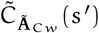 is greater than a_NN_.

