## Supplementary Information for "On biological networks capable of robust adaptation in the presence of uncertainties: A systems-theoretic approach"

### Supporting Information

#### 0.1 Proof of Claim 1

:

*Proof.* Since the external disturbance $d(t)$ has a direct effect on the dynamics of only $x_1$ equation (38) can be rewritten as

$$\dot{x}_1 = \mathbf{A}_{1,1}x_1 + \mathbf{A}_{\tilde{x}}\tilde{x} + b_{1y}y + b_1 d \tag{1}$$

$$\dot{\tilde{\mathbf{x}}} = \tilde{\mathbf{A}}\tilde{\mathbf{x}} + \mathbf{A}_1 x_1 + \mathbf{B}_z \mathbf{z}(t) + \mathbf{B}_y y \tag{2}$$

$$\dot{y} = \alpha_{1,y}x_1 + \mathbf{B}_{\tilde{x}}\tilde{\mathbf{x}} + \mathbf{B}_{z,y}\mathbf{z} + \alpha_{y,y}y \tag{3}$$

$$\dot{\mathbf{z}} = \mathbf{A}_z\mathbf{z} + \mathbf{B}_{1x}x_1 + \tilde{B}_{\tilde{x}}\tilde{\mathbf{x}} + \mathbf{B}_{y,z}y \tag{4}$$

where, $\tilde{x} \in \mathbb{R}^{N-2}$ are the states apart from the input-receiving state and the output state $(y)$. Transforming the set of equations (2)-(4) in Laplace domain we obtain

$$\begin{bmatrix} \mathbf{B}_y & \mathbf{B}_z \\ (s - \alpha_{y,y}) & \mathbf{B}_{z,y} \end{bmatrix} \begin{bmatrix} Y(s) \\ \mathbf{Z}(s) \end{bmatrix} = \begin{bmatrix} s\mathbf{I}_{N-2\times N-2} - \tilde{\mathbf{A}} & \mathbf{A}_1 \\ \mathbf{B}_{\tilde{x}} & \alpha_{1,y} \end{bmatrix} \begin{bmatrix} \tilde{\mathbf{X}}(s) \\ X_1(s) \end{bmatrix} \tag{5}$$

Since, the disturbance $d$ is only applied to the node $x_1$, the controllability condition for the pair $(\mathbf{A}_w, \mathbf{E}_w)$ translates to the same with the pair $(\tilde{\mathbf{A}}, \mathbf{A}_1)$. Therefore the matrix $\begin{bmatrix} s\mathbf{I}_{N-2\times N-2} - \tilde{\mathbf{A}} & \mathbf{A}_1 \end{bmatrix}$ is always structurally full rank (P-B-H condition for controllability). The only way the system of linear equations in equation (5) shall have non-unique or no solution if the last row of the co-efficient matrix of $\tilde{\mathbf{X}}(s)$ and $X_1(s)$ is zero which is in contradiction with the output controllability assumption. Therefore, in case of full controllability, $\tilde{\mathbf{X}}(s)$ and $X_1(s)$ can always be expressed in terms of $\tilde{\mathbf{Z}}(s)$ and $Y(s)$. Further, using equation (4), it is always possible to write $Z(s)$ in the desired form □

**Remark 3.** *The strict assumption of full structural controllability of the pair* $(\mathbf{A}_w, \mathbf{E}_w)$ *can be relaxed further. Let us assume that there exists a module including* $K$ *interconnected nodes that is not controllable by node* $x_1$ *but is influenced by the control module* $\mathbf{z}$*. Therefore, the corresponding Laplace equations can be written as*

$$\begin{bmatrix} \mathbf{B}_{y,K} & \mathbf{B}_{z,K} \\ \mathbf{B}_{y,N-2-K} & \mathbf{B}_{z,K} \\ (s - \alpha_{y,y}) & \mathbf{B}_{z,y} \end{bmatrix} \begin{bmatrix} Y(s) \\ \mathbf{Z}(s) \end{bmatrix} = \begin{bmatrix} s\mathbf{I}_{K\times K} - \tilde{\mathbf{A}}_K & \mathbf{0}_{K\times N-K-2} & \mathbf{0}_K \\ \tilde{\mathbf{A}}_{K,N-K-2} & \tilde{\mathbf{A}}_{N-K-2,N-K-2} & \mathbf{A}_{1,N-K-2} \\ \mathbf{B}_{K,\tilde{x}} & \mathbf{B}_{N-K-2,\tilde{x}} & \alpha_{1,y} \end{bmatrix} \begin{bmatrix} \tilde{\mathbf{X}}_K(s) \\ \tilde{\mathbf{X}}_{N-K-2}(s) \\ X_1(s) \end{bmatrix} \tag{6}$$

*From the equation* (6)*, it can be seen that* $\tilde{\mathbf{X}}_K(s)$ *can be solved from the first* $K$ *equations. Therefore, the solution* $[\tilde{\mathbf{X}}_K(s)\ \tilde{\mathbf{X}}_{N-K-2}(s)\ X_1(s)]$ *is unique if and only if there exists at least one element in* $\mathbf{B}_{K-N-2,\tilde{x}}$ *or* $\alpha_{31}$ *is non-zero—this translates the the claim 1 can be made possible even when the network is not fully controllable by the external disturbance* $d$ *given the output is controllable by the same and the 'uncontrollable-by-disturbance' modules are controllable by the proposed control input.*

#### 0.2 Proof for Proposition 1

*Proof.* In the first case, $A_z = 0$, therefore the system matrix for the augmented state space $\mathbf{x}_c := [\mathbf{x}\ z]'$ in the closed-loop system

$$\mathbf{A}_{Cw} = \begin{bmatrix} \mathbf{A}_w & \mathbf{B}_w \\ \mathbf{B}_x & A_z \end{bmatrix}$$

Since perfect adaptation implies stability, the matrix $\mathbf{A}_{Cw}$ has to be Hurwitz. This translates to a set necessary conditions that require all the coefficients of $s^k$, $\forall k = 0(i)N-1$ of the monic characteristics polynomial $P_{\mathbf{A}_{Cw}}(s) := \mathrm{Det}(s\mathbf{I}_{N\times N} - \mathbf{A})$ to be positive. Therefore, the determinant condition for stability can be written as

$$\mathrm{sign}(\mathrm{Det}(\mathbf{A}_{Cw})) = (-1)^N$$

From combinatorial matrix theory, each of the N! terms in the determinant expression of $\mathbf{A}_{Cw}$ contains a multiplicative combination of either the diagonals or the combination of off-diagonal elements in such a way that these off-diagonal combinations map to a loop (cycle) in the actual network structure [39]. Therefore, in the case where one of the diagonal elements ($A_z$) of the closed-loop system matrix $\mathbf{A}_{Cw}$ is zero, all the terms in the determinant expression of $\mathbf{A}_{Cw}$ contain at least one loop. Further, Bhattacharya *et al* (2022) proved that in the case of buffer action (one or more diagonal elements of $\mathbf{A}_{Cw}$ is/are zero), the network has to contain at least one negative cycle to achieve stability. This concludes the first part of the proof [27].
Further, as the condition suggests, only the negative feedback with buffer action (*i. e.* $A_z = 0$) is not sufficient for the mentioned network structure and does not guarantee a zero $\tilde{G}_1(s)$ that is crucial for producing an integral control action. $\tilde{G}_1(s)$ can be made zero if the (concentration of the) set of process nodes having a direct edge to the controller (denote as $\mathcal{N}_{pz}$) can be expressed as a sole function of the output node ($Y(s)$). This, in turn, is possible in an P−node, P− edge network, if there exists no incoming edge from the controller either to any of the paths from the process nodes in $\mathcal{N}_{pz}$ to the output node or from the output to the process nodes in $\mathcal{N}_{pz}$. Fig. 4 illustrates the claim. □

#### 0.3 Proof of Proposition 2

*Proof.* For the closed-loop system, the transfer function between the output ($Y(s)$) and the disturbance signal ($D(s)$) can be written as

$$\frac{Y(s)}{D(s)} = \mathbf{C}(s\mathbf{I}_{N\times N} - \mathbf{A}_{Cw})^{-1}\mathbf{E}_w$$

Let us denote the concentration of the $K^{th}$ node as the output variable. Because the $\mathbf{E}_w$ is a unit vector along the direction of $x_1$, it can be shown that the numerator polynomial of the transfer function can be derived just by computing the minor of the $(1,k)$ element of $(s\mathbf{I}_{N\times N} - \mathbf{A}_{Cw})^{-1}$. The denominator is the characteristics polynomial of $\mathbf{A}_{Cw}$ in terms of the Laplace variable s. Similarly, the transfer function between the controller node ($Z(s)$) (denote as the $j^{th}$ node) and the disturbance input

$$\frac{Z(s)}{D(s)} = \tilde{\mathbf{C}}(s\mathbf{I}_{N\times N} - \mathbf{A}_{Cw})^{-1}\mathbf{E}_w \tag{7}$$

$$\frac{Z(s)}{Y(s)} = \frac{\tilde{\mathbf{C}}(s\mathbf{I}_{N\times N} - \mathbf{A}_{Cw})^{-1}\mathbf{E}_w}{\mathbf{C}(s\mathbf{I}_{N\times N} - \mathbf{A}_{Cw})^{-1}\mathbf{E}_w} \tag{8}$$

$$\implies \frac{\mathrm{Minor}\Big(\big[s\mathbf{I}_{N\times N} - \mathbf{A}_{Cw}\big]_{1,j}\Big)}{\mathrm{Minor}\Big(\big[s\mathbf{I}_{N\times N} - \mathbf{A}_{Cw}\big]_{1,k}\Big)} \tag{9}$$

It is worth noting that the coefficient of $s^0$ in the denominator coefficient is the minor of $\big[\mathbf{A}_{Cw}\big]_{1,k}$. For the controller to provide the integral action, the denominator polynomial should be divisible by s—this translates to the requirement of the $M =$

$\text{Minor}\left([\mathbf{A}_{Cw}]_{1,k}\right)$ to be zero. Further, the quantity $\left[\mathbf{A}_{Cw}\right]_{1,k} \times M$ is an element in the determinant expression of $\mathbf{A}$. Therefore, the quantity $M$ encodes all possible forward paths, multiplied by the diagonal elements, from the node $x_1$ to the output node $x_k$. Now, in the first case, when $A_z = 0$ all the elements of $M$ containing the diagonal element $A_z$ go to zero. Therefore, through a further choice of edges, it becomes feasible to make all the elements in $M$ individually zero without making the system structurally uncontrollable. Unlike in the first case, if $A_z \neq 0$, there exist multiple elements in $M$. Following this, two forward paths from the input-receiving to the output nodes with opposing effects (the diagonals are assumed to be non-positive). This concludes the proof. Figure 5 illustrates the claim. □

#### 0.4 Proof for Proposition 3

*Proof.* An opposer module, as defined in the previous sections, does not contain any cycle—this implies that the eigenvalues of the system matrix associated with the opposer modules are the diagonal elements. Therefore, the corresponding characteristics equation of $\mathbf{A}_{Cw}$ in case of an opposer module can be

$$C_{\mathbf{A}_{Cw}}(s) = s^N + \sum_{k=1}^{N}(-1)^k \sum \sigma_k(\rho(\mathbf{A}_{Cw}))s^{N-k}$$

where, $\sigma_k$ is a k-length permutation operator, $\sigma_k(\rho(\mathbf{A}))$ returns the set containing all possible $k$–length multiplicative combinations of diagonals from the spectrum of $\mathbf{A}_{Cw}$ $(\rho(\mathbf{A}_{Cw}))$. Evidently, $\sum \sigma_k(\rho(\mathbf{A}_{Cw}))s^{N-k}$ refers the sum across the the set $\sigma_k$.
After addition of a negative feedback loop involving P nodes, the corresponding characteristics polynomial of the modified closed-loop matrix $\tilde{\mathbf{A}}_{Cw}$ can be written as

$$\tilde{C}_{\tilde{\mathbf{A}}_{Cw}}(s) = s^N + \sum_{k=1}^{N}(-1)^k \sum \sigma_k(\rho(\mathbf{A}_{Cw}))s^{N-k} + \sum_{k=P}^{N}(-1)^{(k+P-1)}\mathcal{L}_P \sum(\sigma_{k-P}(\rho(\mathbf{A}_{Cw}))|\mathcal{N}(\mathcal{L}_P))s^{N-k}$$

where, $\mathcal{L}_P$ encodes the P-node loops, $\mathcal{N}(\mathcal{L}_P)$ returns the nodes (rows of $\tilde{\mathbf{A}}_{Cw}$) involved in $\mathcal{L}_P$ $\sigma_{k-P}(\rho(\mathbf{A}_{Cw}))|\mathcal{N}(\mathcal{L}_P)$ returns the set of $k-P$ length multiplicative combination of diagonals of the rows other than the ones occupied by the loop. Since $\text{sign}(L)_P = -1$, the term $(-1)^{k+P-1}\mathcal{L}_P \sum(\sigma_{k-P}(\rho(\mathbf{A}_{Cw}))|$ is always positive. Without any loss of generality, we assume that the diagonal $a_{NN}$ is of the smallest magnitude among the spectrum of $\mathbf{A}_{Cw}$ (Note, all the diagonals are by design negative). Therefore, using the concept of relative stability we can draw valuable insights about the change in the the pole positions after addition of the loop. Replacing $s = s' + a_{NN}$ in the modified characteristic equation

$$\tilde{C}_{\tilde{\mathbf{A}}_{Cw}}(s') = s^N + \sum_{k=1}^{N}(-1)^k \sum \sigma_k(\rho(\mathbf{A}_{Cw}))(s' + a_{NN})^{N-k} + \sum_{k=P}^{N}(-1)^{(k+P-1)}\mathcal{L}_P \sum(\sigma_{k-P}(\rho(\mathbf{A}_{Cw}))|\mathcal{N}(\mathcal{L}_P))(s' + a_{NN})^{N-k}$$

Further, through appropriate grouping of the terms (Supporting information) $\tilde{C}_{\tilde{\mathbf{A}}_{Cw}}(s')$

can be expressed as

$$\tilde{C}_{\tilde{\mathbf{A}}_{Cw}}(s') = s^N + \sum_{k=1}^{N}(-1)^k \sum \sigma_k(\rho(\mathbf{A}_{Cw} - a_{NN}))s'^{N-k} + (-1)^N \mathrm{Det}(\tilde{\mathbf{A}}_{Cw}) + \sum_{k=P}^{N-1}(-1)^{(k+P-1)}\mathcal{L}_P \sum (\sigma_{k-P}(\rho(\mathbf{A}_{Cw} - a_{NN}))|\mathcal{N}(\mathcal{L}_P))(s')^{N-k}$$

where $\rho(\mathbf{A}_{Cw} - a_{NN})$ returns the set of all the diagonals of $\mathbf{A}_{Cw}$ (except $a_{NN}$) deducted by an amount of $a_{NN}$. Since, $|a_{NN}|$ is the minimum across the diagonals of $\tilde{\mathbf{A}}_{Cw}$ the quantity $a_{ii} - a_{NN}$ remains negative where $a_{ii}$ is any other diagonal of $\tilde{\mathbf{A}}_{Cw}$. Further, because of the negative feedback the coefficient of $s'^0$ remains positive. In fact, it can be shown that $\mathrm{Det}(\tilde{\mathbf{A}}_{Cw}) > \mathrm{Det}(\mathbf{A}_{Cw})$. Therefore, since all the coefficients of $\tilde{C}_{\tilde{\mathbf{A}}_{Cw}}(s')$ are positive, it can be shown that the magnitude of the closest root of $\tilde{C}_{\tilde{\mathbf{A}}_{Cw}}(s')$ is greater than $a_{NN}$. □
